# Tracking human population structure through time from whole genome sequences

**DOI:** 10.1101/585265

**Authors:** Ke Wang, Iain Mathieson, Jared O’Connell, Stephan Schiffels

## Abstract

The genetic diversity of humans, like many species, has been shaped by a complex pattern of population separations followed by isolation and subsequent admixture. This pattern, reaching at least as far back as the appearance of our species in the paleontological record, has left its traces in our genomes. Reconstructing a population’s history from these traces is a challenging problem. Here we present a novel approach based on the Multiple Sequentially Markovian Coalescent (MSMC) to analyse the population separation history. Our approach, called MSMC-IM, uses an improved implementation of the MSMC (MSMC2) to estimate coalescence rates within and across pairs of populations, and then fits a continuous Isolation-Migration model to these rates to obtain a time-dependent estimate of gene flow. We show, using simulations, that our method can identify complex demographic scenarios involving post-split admixture or archaic introgression. We apply MSMC-IM to whole genome sequences from 15 worldwide populations, tracking the process of human genetic diversification. We detect traces of extremely deep ancestry between some African populations, with around 1% of ancestry dating to divergences older than a million years ago.

**Author Summary:** Human demographic history is reflected in specific patterns of shared mutations between the genomes from different populations. Here we aim to unravel this pattern to infer population structure through time with a new approach, called MSMC-IM. Based on estimates of coalescence rates within and across populations, MSMC-IM fits a time-dependent migration model to the pairwise rate of coalescences. We implemented this approach as an extension to existing software (MSMC2), and tested it with simulations exhibiting different histories of admixture and gene flow. We then applied it to the genomes from 15 worldwide populations to reveal their pairwise separation history ranging from a few thousand up to several million years ago. Among other results, we find evidence for remarkably deep population structure in some African population pairs, suggesting that deep ancestry dating to one million years ago and older is still present in human populations in small amounts today.

## Introduction

Genomes harbour rich information about population history, encoded in patterns of mutations and recombinations. Extracting that information is challenging, since in principle it requires reconstructing thousands of gene genealogies separated by ancestral recombination events, using only the observable pattern of shared and private mutations along multiple sequences. One important innovation was the Sequentially Markovian Coalescent (SMC) model[1,2], which is an approximate form of the ancestral recombination graph that can be fitted as a Hidden Markov model along the sequence. This approach has been used to infer demographic history in methods like PSMC[3], MSMC[4], diCal[5,6] and SMC++[7].

These methods estimate one or both of two important aspects of population history: i) The history of the effective population size, and ii) the history of population structure. The second aspect, which entails reconstructing the timing and dynamics of population separation requires a non-trivial choice of parameterization: While methods like diCal2[5], as well as many methods based on the joint site frequency spectrum[8–11] use an explicit population model with split times, migration rates or admixture events, MSMC [4] introduced the concept of the relative cross coalescence rate to capture population separations in a continuously parameterised fashion. The main advantage of that approach is that it does not require the specification of an explicit model, but can be applied hypothesis-free to estimate key aspects of population separation, for example the time at which lineages are half as likely to coalesce between rather than within populations, which is often used as a heuristic estimate for the divergence time between the populations. A disadvantage is that other important aspects of population separation, like post-split or archaic admixture, are non-trivially encoded in features of the cross-coalescence rate other than this mid-point. As a consequence, it is difficult to interpret the cross-coalescence rate in terms of actual historical events.

Here, we propose an approach to overcome the disadvantages of the relative cross coalescence rate, while maintaining the continuous character of population separation from MSMC without explicitly specifying a complex population phylogeny. We present a new method MSMC-IM, which fits a continuous Isolation-Migration (IM) model to the distribution of coalescence times, estimated from MSMC’s piecewise constant model. In MSMC-IM, separation and migration between a pair of populations is quantified by a piecewise constant migration rate across populations, and piecewise constant population size changes within each population. We apply our method on world-wide human genomic data from the Simons Genome Diversity Project (SGDP)[12] to investigate the history of global human population structure.

## Results

### Estimating pairwise coalescence rates with MSMC2 and fitting an IM model

To model the ancestral relationship between a pair of populations, we developed an isolation-migration model with a time-dependent migration rate between a pair of populations, which we call MSMC-IM. The approach requires time-dependent estimates of pairwise coalescence rates within and across two populations. Here we use MSMC2 for these estimates, which was first introduced in Malaspinas et al. 2016[13] (see Methods). MSMC2 offers two key advantages over MSMC[4]. First, the pairwise coalescence model in MSMC2 is exact within the SMC’ framework[2], whereas MSMC’s model uses approximations that cause biases in rate estimates for larger number of haplotypes (S1 Fig). Second, since MSMC2 uses the pairwise tMRCA distribution instead of the first tMRCA distribution, it estimates coalescence rates within the entire range of coalescence events between multiple haplotypes, which ultimately increases resolution not just in recent times but also in the deep past. These two improvements are crucial for our new method MSMC-IM, which relies on unbiased coalescence rate estimates within and across populations, in particular in the deep past. Specifically, MSMC2 recovers simulated population size histories (with human-like parameters) well up to 3 million years ago, while keeping the same high resolution in recent times as MSMC (S1 Fig).

Given MSMC2’s estimates of time-dependent coalescence rates within populations, λ_11_(t) and λ_22_(t), and across populations, λ_12_(t), we use MSMC-IM to fit an Isolation-Migration (IM) model to those three coalescence rates (see Methods). MSMC-IM’s model assumes two populations, each with its own population size *N*_1_(*t*) and N_2_(t), and a piecewise-constant symmetric migration rate *m(t)* between the two populations (Fig 1B, see Methods and S1 Text for details). Expressing the separation history between two populations in terms of a variable migration rate instead of the more heuristic relative cross coalescence rate facilitates interpretation, while maintaining the freedom to analyse data without having to specify an explicit model of splits and subsequent gene flow.

**Fig 1.**
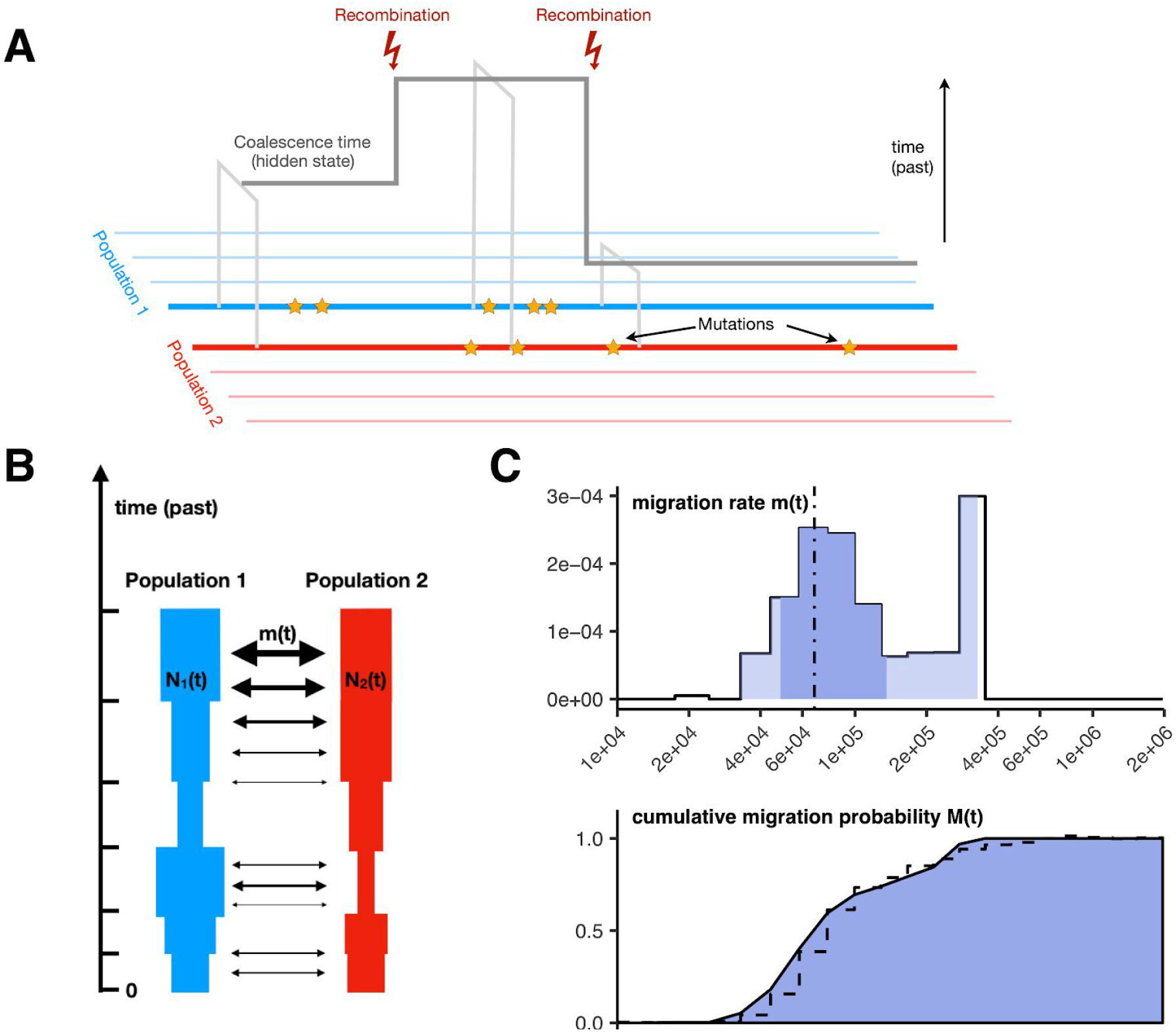
Schematic of MSMC2 and MSMC-IM. (A) MSMC2 analyses patterns of mutations between pairs of haplotypes to estimate coalescence local times along the genome. (B) MSMC-IM fits an isolation-migration model to the pairwise coalescence rate estimates, with time-dependent population sizes and migration rate. (C) As a result, we obtain migration rate densities and cumulative migration probabilities for pairs of populations.

### Evaluating MSMC-IM with simulated data

We illustrate MSMC-IM by applying to several series of simulated scenarios of population separation (see Methods). First, the *clean-split*-scenario consists of an ancestral population that splits into two subpopulations at time T (Fig 2A). Second, the *split-with-migration*-scenario adds an additional phase of bidirectional gene flow between the populations after they have split (Fig 2B). Third, the *split-with-archaic-admixture*-scenario involves no post-split gene flow, but contains additional admixture into one of the two extant populations from an unsampled “ghost” population, which splits from the ancestral population (Fig 2C) at time *T_a_*>*T*. For each scenario, we simulated 8 haplotypes (four from each population), used human-like evolutionary parameters and varied one key parameter to create a series of related scenarios (see Methods). We then ran MSMC2 to estimate coalescence rates for haplotype pairs within each population and across populations, and applied MSMC-IM to fit our IM model with variable migration to these coalescence rates.

**Fig 2.**
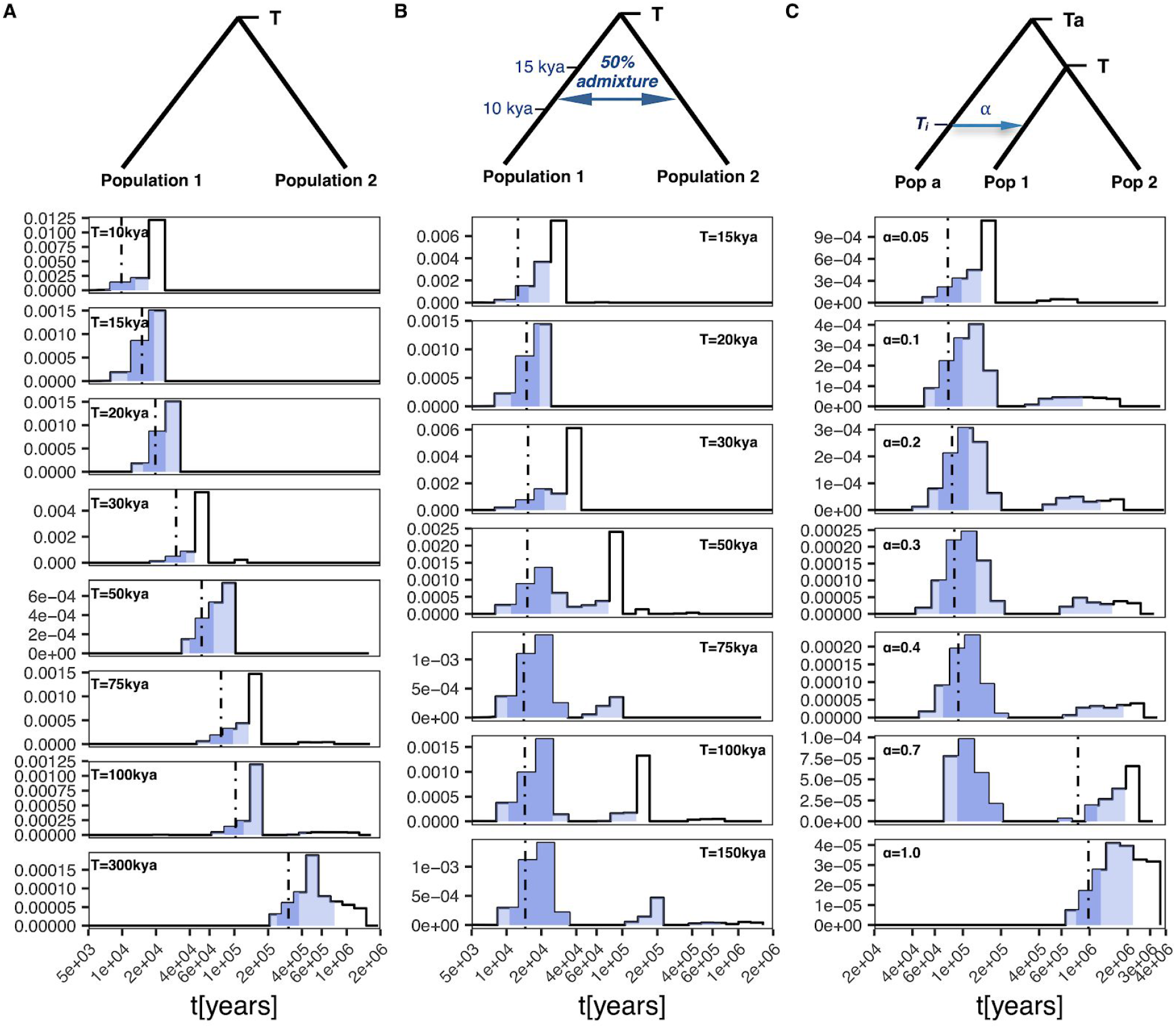
Simulation results. **(A) *Clean-split-scenario***. We simulated 8 haplotypes intotal from two populations with constant size 20,000 diverged at split time *T* varying from 10 thousand years ago (kya) to 300kya. **(B) *Split-with-migration*-scenario**. Similar to A), with *T* varying between 15-150kya, and, with a time period of symmetric migration between 10kya to 15kya. The cumulative migration rate is tuned such that 50% of lineages migrate in total. **(C) *Split-with-archaic-admixture*-scenario**. Similar to A), with *T*=75kya. Here, population 1 receives an admixture pulse at 30kya from an archaic ghost population, which separates from the ancestral population at 1mya. The admixture rate varies from 5% to 100%. In all plots, the blue light blue shading indicates the interval between 1-99% of the cumulative migration probability, the dark blue shade from 25-75%, and the vertical line indicates the median.

In the clean-split scenario, we find that the inferred migration rate *m(t)* displays a single pulse of migration around the simulated split time *T* (Fig 2A). This is expected, since in our parametrization, a population split corresponds to an instantaneous migration of lineages into one population at time *T*, thereby resulting in a single pulse of migration. In the *split-with-migration* series, we expect two instead of one pulse of migration: one at time *T*, as above, and a second more recent one around the time of post-split migration. In cases where the split time and migration phase are separated by more than around 20,000 years, this is indeed what we see (Fig 2B), although with some noise around this basic pattern. For less time of separation of the two migration pulses, MSMC-IM is not able to separate them in this scenario. Similarly, we find two phases of migration for the *split-with-archaic-admixture*-scenario, with one phase around time *T*, and another one around the time of the archaic split (Fig 2C).

Viewing these results in a different way, we can use the estimated migration rate *m(t)* to compute the probability that a lineage sampled at present has migrated into the other subpopulation, looking backwards in time. Specifically, we define the probability

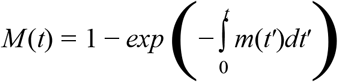

which continuously increases from 0 to 1 in all scenarios, with gradient zero at times of no migration, and strictly positive gradient in periods of migration (S2 Fig). Numerically, it turns out that *M(t)* is similar to the relative cross coalescence rate (CCR), thereby heuristically justifying the CCR as a measure of population separation. When *M(t)* is very close to 1, enough migration has occurred that lineages sampled at present in one of the two subpopulations have lost their subpopulation identity at that point in the past, and in fact the two populations in the IM model can be seen as one from this time towards the past. In theory, when *M(t)* becomes very close to 1, our three-parameter model is overspecified, and there should be only a single parameter left, the ancestral population size. We solve this problem by regularization (Methods)–penalising differences between the two population sizes, and by ignoring migration rate estimates at times when *M(t)* >0.999.

MSMC-IM also fits population sizes, which can be compared to the raw estimates from MSMC, i.e. to the inverse coalescence rates within population 1 and 2, respectively (see S1 Text for some non-trivial details on this comparison). We find that estimates for *N_1_(t)* and *N_2_(t)* are in fact close to the inverse coalescence rates, without much effect from the estimated migration rates (S3 Fig).

### Deep ancestry in Africa

We applied our model to 30 high coverage genomes from 15 world-wide populations from the SGDP dataset[12] (S1 Table) to analyse global divergence processes in the human past. When analysing the resulting pairwise migration rate profiles, we find that several population pairs from Africa exhibit by far the oldest population structure observed in all pairwise analyses. We find that in all population pairs involving either San or Mbuti, the main separation process from other populations dates to between 100-400 thousand years ago, depending on the exact pair of populations (see below), but with small amounts reaching back to beyond a million years ago, as seen by the non-zero migration rates around that time (Fig 3A, S4 Fig), and the cumulative migration probability, *M(t)*, (Fig 3B) which has not fully reached 1 until beyond a million years ago. The genetic separation profile in pairs involving Mbuti and San is, beyond the extraordinary time depth, not compatible with clean population splits (as seen in simulations, Fig 2A) or simple scenarios of archaic admixture, but instead shows evidence for multiple periods of gene flow between (unsampled) populations. Between Mbuti and other African populations except San, we find three distinct phases of gene flow. The first peaks around 15,000 years ago, compatible with relatively recent admixture between Mbuti and other African populations. The second phase spans from 60 to 300 thousand years ago, reflecting the main genetic separation process, which itself looks complex and exhibits two peaks around 80 and 200 thousand years ago. The third and final phase, including a few percent of lineages from around 600,000 to 2 million years ago, likely reflects admixture between populations that diverged from each other at least 600,000 years ago. In pairs that include San, the onset of gene flow with other populations is more ancient than with Mbuti, beginning at around 40,000 years ago and spanning until around 400,000 years ago in the main phase, and then exhibiting a similarly deep phase as seen in Mbuti between 600,000 and 2 million years ago. We confirm that this deep divergence is robust to phasing strategy (see below) and filtering (see Methods). We also replicated this signal using an independent dataset[14] (S5 Fig).

**Fig 3.**
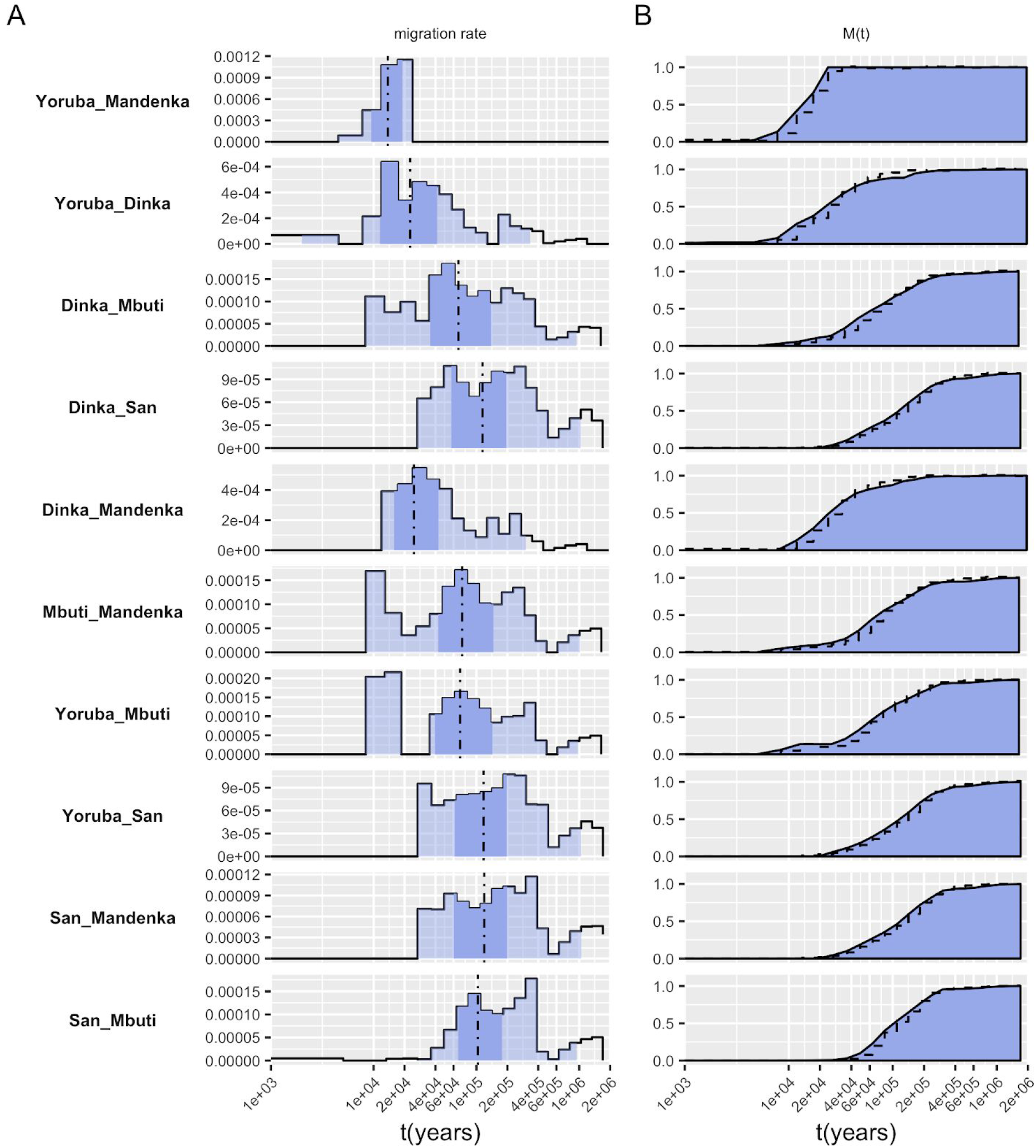
Migration structure within 5 African populations. (A) Migration rates. (B) Cumulative migration rates M(t). Color shading as in Fig. 2.

Apart from the deep structure seen with Mbuti and San, we find similarly deep signals between the West African Yoruba, Mandenka and Mende on the one hand, and the East African Dinka on the other (Fig 3A, S4 Fig A-F), suggesting archaic ancestry with differential relationship between West and East Africans. This might be consistent with recent findings of archaic ancestry in West-Africans[15,16], or with West-Eurasian ancestry in East Africans[17], which may carry Neanderthal ancestry. Finally, pairwise analyses among Mende, Mandenka and Yoruba (Fig 3A, S4 Fig C,E,F) exhibit a very recent migration profile, which appears to span up to about 20,000 years ago but not older, which is at odds with a recent finding of basal African ancestry present to different degrees in Mende and Yoruba [18]. However, that signal may be too weak to be detected in our method, which is based on only two individuals per population. Also, the basal African ancestry detected in [18] was inferred to be younger than the split of Neanderthal and Denisovan from the main human lineage, and therefore might be too close to the main phase of population differentiation in Africa to be detected by MSMC-IM (see simulation results above).

### Complex Out-of-Africa

In contrast to the variety of separation profiles between African populations, most profiles between African and Non-African populations look remarkably similar, with a main separation phase between 40 and 150 thousand years ago, and a separate peak between 200 and 400 thousand years ago (Fig 4). The first, more recent, phase plausibly reflects the main separation of Non-African lineages from African lineages predating the “out-of-Africa” migration event, with signals more recent than about 60,000 years likely reflecting the typical spread of MSMC-estimated coalescence rate changes observed previously [4]. The second peak of migration between 200 and 400 thousand years ago likely reflects Neandertal and/or Denisovan introgression into non-Africans. The age of that peak appears too recent given previous split time estimates of those two Archaic groups from the main human lineage at 550-765 thousand years ago (kya)[14]. However, our simulation with archaic admixture (Fig 2C), shows that our model tends to underestimate the archaic split time when the migration proportion is low, like the estimated proportions of Neanderthal and Denisovan ancestry in modern humans, which are both below 10%[19–21].

**Fig 4.**
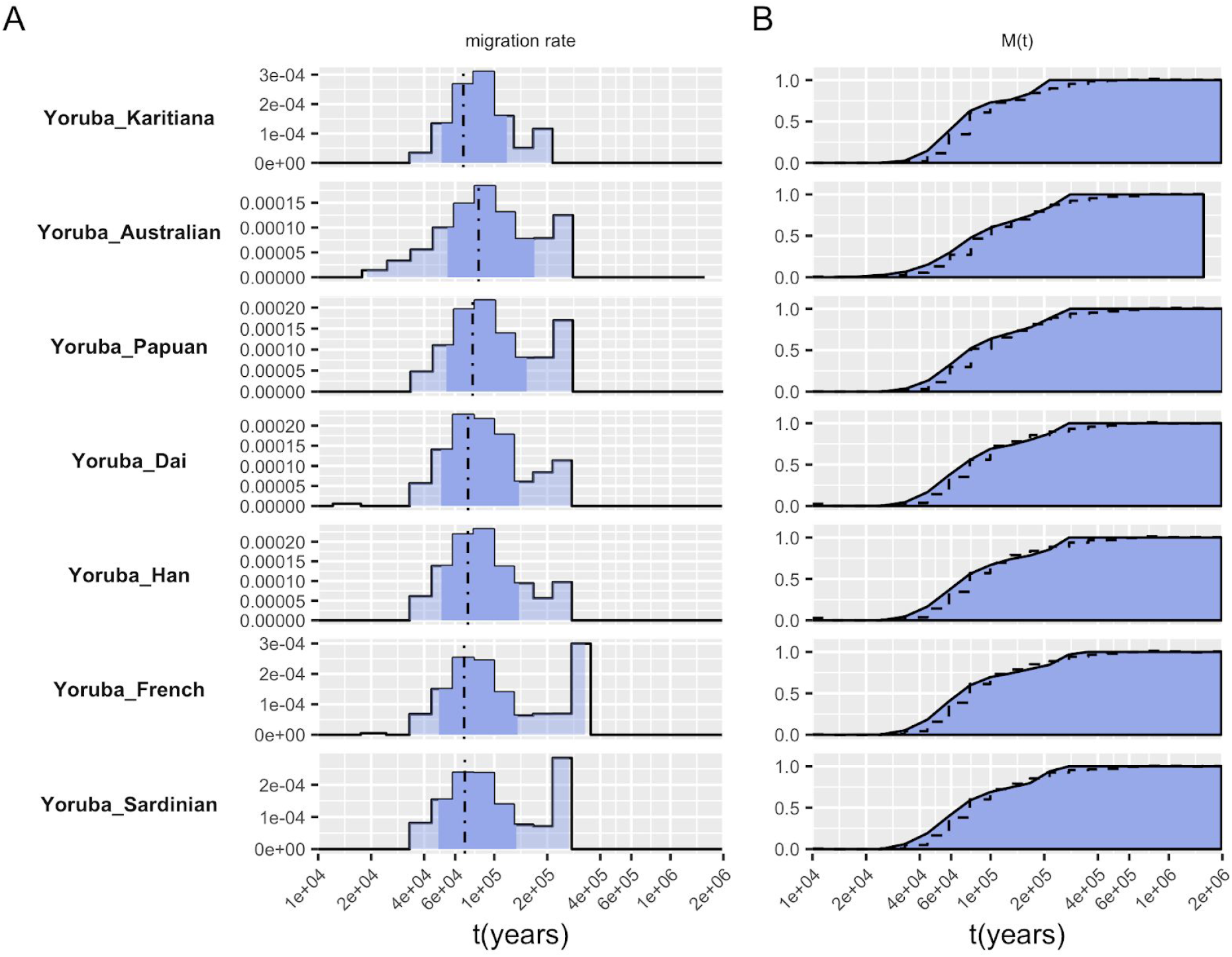
Selected migration profiles between Yoruba and 7 non-African populations. (A) Migration rates. (B) Cumulative migration rates M(t).

The majority of African/Non-African population pairs follows this simple pattern, but there are some exceptions that exhibit a deeper migration profile, more similar to that seen within Africa (S4 Fig). In particular, French exhibit deep migration structure with San, Mbuti and Mandenka, and similar signals are seen with Karitiana (with Mandenka), Sardinian (with San) and Han (with Mandenka). In principle, since Non-African populations descend from an African population, all African/Non-African migration profiles should in turn exhibit a similarly deep migration profile with San, Mbuti and perhaps some West African populations as we have seen in African pairs. That we do not see such deep structure in most Non-African/African pairs is likely due to the strong genetic drift that the Non-African ancestors experienced during the Out-of-Africa bottleneck, as documented in the marked population size decline around 60kya, seen in MSMC2 population size plots (S6 Fig).

We investigated previous observations of potential ancestry from an earlier dispersal out of Africa, present in Papuan and Australian genomes[12,13,22]. While we were able to replicate the slight shift of rCCR or M(t) midpoint-based split times from African/Eurasian pairs to African/Australasian pairs reported in Ref.[22] using MSMC and Ref.[13] using MSMC2, we find that the estimated migration profiles of these pairs are very similar (S7 Fig), with a main separation mid-point around 70kya and a second older signal beyond 200kya, consistent with both Australasians and other Non-Africans being derived from a single genetic ancestral population without a more basal contribution to Australasians[12,13]. We note, however, that different separation events are not distinguishable in MSMC-IM when they are temporarily close to each other, as we have seen in the *split-with-migration-scenario* (Fig. 2B).

### Separations outside of Africa

All separations outside of Africa are younger than separations between Africans and Non-Africans, as expected (Fig 5, S4 Fig G-M). The deepest splits outside of Africa are seen in pairs of Papuans or Australians with other Eurasians, in which the first peak of migration is seen at 40-60 kya, corresponding to the early separation of these populations’ ancestors from other non-African populations after the out of Africa dispersal. In these pairs we see a second peak around 250-300 kya, likely corresponding to the known Denisovan admixture in Papuans and Australians[13,23]. As discussed above, this is too recent for divergence time estimates between Denisovans and modern humans[14], which again is likely due to the relatively low levels of admixture, which we showed in simulations can lead to underestimates of the divergence time. Surprisingly, we see a similar second peak between French and Han, which is consistent with previous observations[4,12] but of unclear cause. Consistent with the hypothesis that the second peak seen in Australasian/Eurasian pairs corresponds to Denisovan admixture, we do not see a second peak in the migration profile between Papuans and Australians, confirming that the gene flow likely occurred into the common ancestor of Australians and Papuans[13]. The migration profile between Papuans and Australians shows a main separation between 15-35 kya.

**Fig 5.**
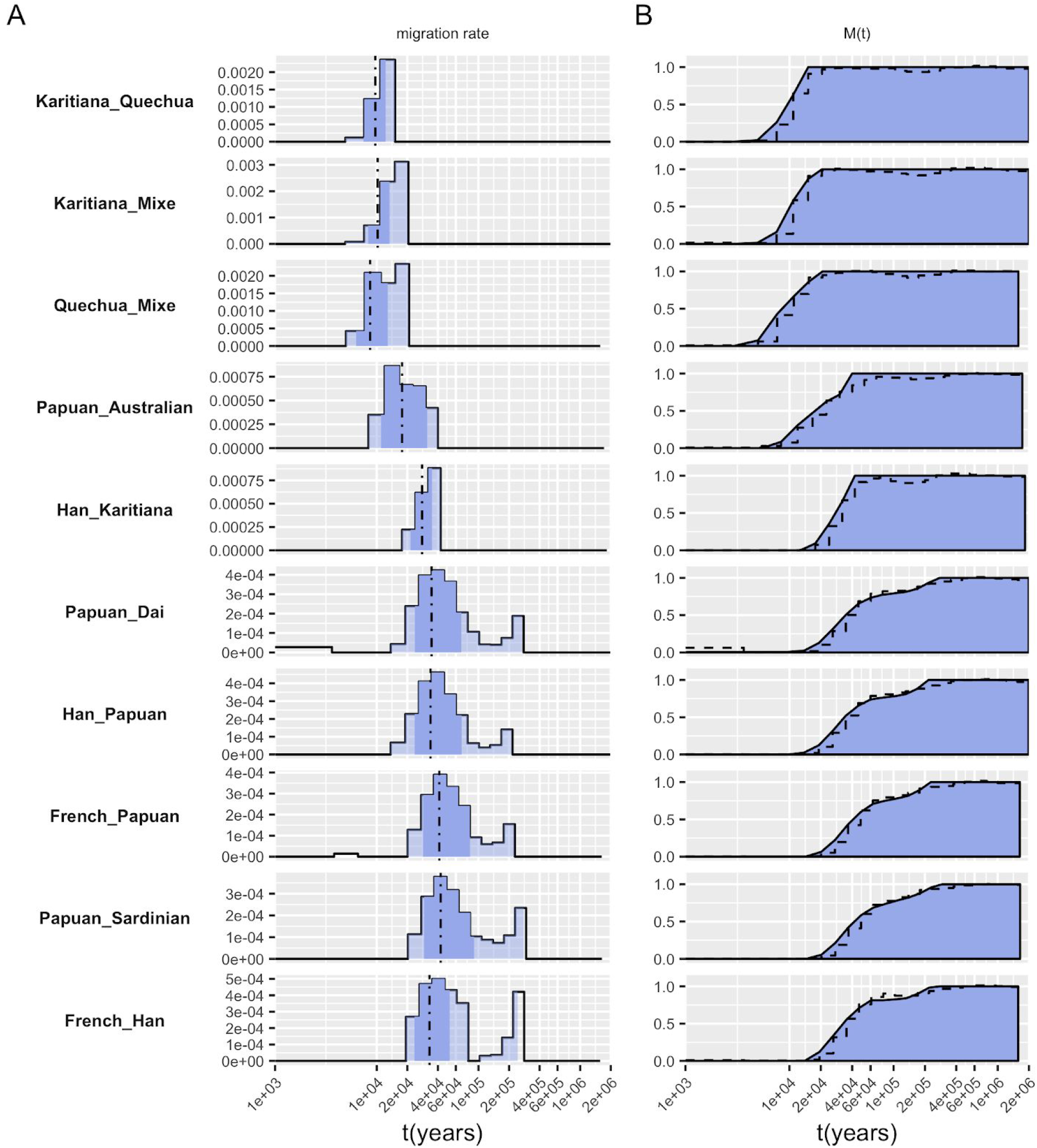
Migration structure within non-African populations. (A) Migration rates. (B) Cumulative migration densities M(t).

The second deepest splits in Non-African populations are seen between East Asian and European populations, which occur mostly between 20 and 60kya (cumulative migration density midpoint at 34kya), followed by separations between Asian and American populations, between 20 and 40kya (midpoint at 26kya). The latter likely also reflects Ancestral North Eurasian ancestry in Americans[24], which is more closely related to Europeans than to East Asians, thereby pushing back the separation seen between East Asians and Native Americans. Finally, the most recent splits are seen between populations from the same continent: Dai/Han split around 10-40kya (midpoint 11kya), French/Sardinian around 8-25kya (midpoint 9kya) and within Native Americans around 7-11kya (midpoint 9.5kya) (Fig 5, S4 Fig).

To visualise the depth of ancestry in each population pair, we summarised all pairwise analyses by percentiles of the cumulative migration density *M(t)* (Fig 6). Largely, Non-African pairs (orange) have their main separation phase, with the cumulative migration density between 25% and 75%, between 20 and 60kya, with some more recently diverged pairs within continents. In contrast, African pairs (red) have their main phase largely between 60 and 200kya, with some notable exceptions of more recently diverged populations, and with the notable tail (99% percentile) up to 1 million years and older. Between Africans and Non-Africans, divergence main phases are largely within a similar window of 60-200kya as in African pairs, with three notable groups: divergence of Non-Africans from San falls between 80-250kya, from Mbuti between 70-200kya, and from other Africans between 50-150kya.

**Fig 6.**
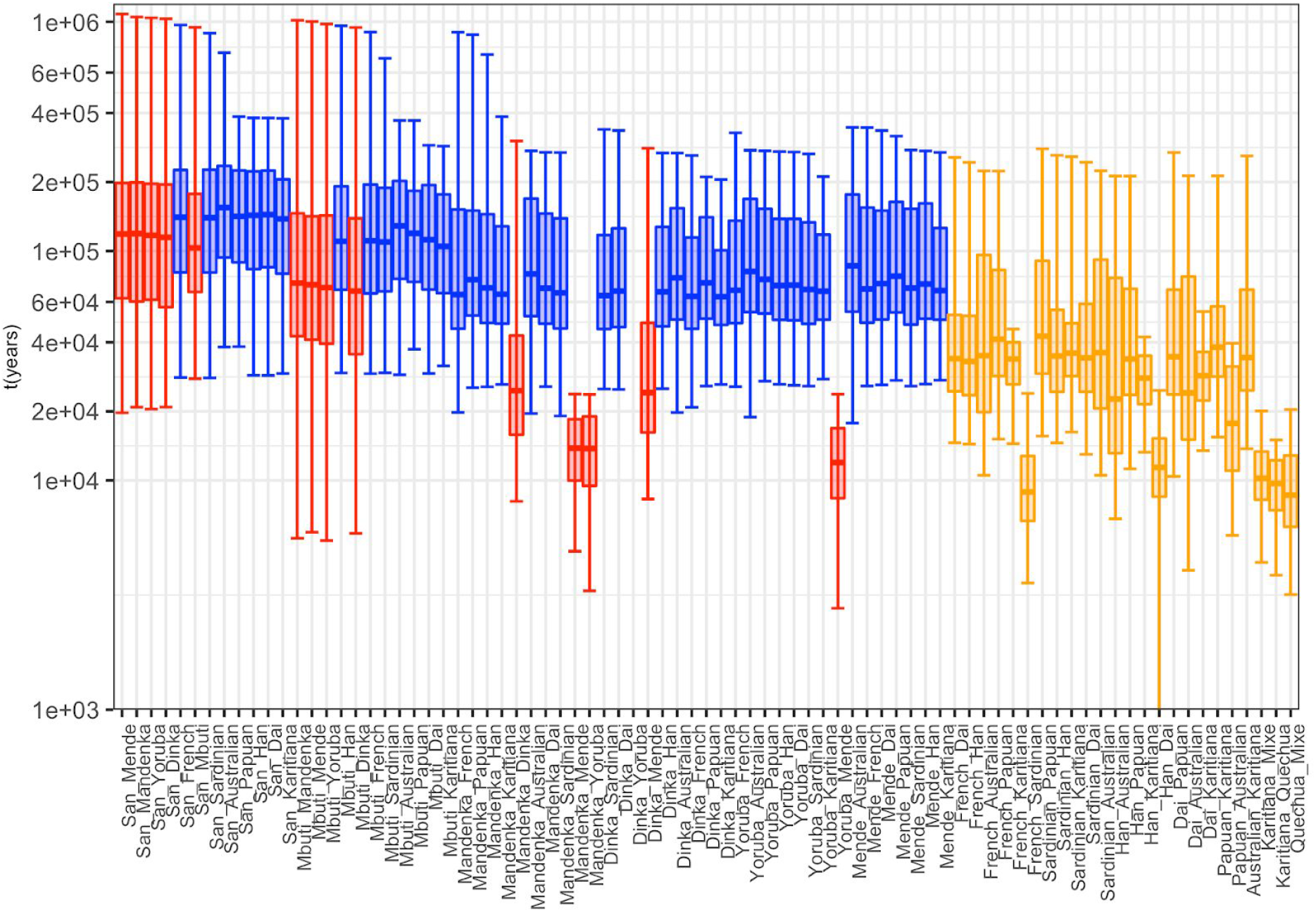
Overview of *M(t)* in quantiles for 81 pairs from 15 world-wide population. Boxes show the 25% to 75% quantiles of *M(t)*, with bi-directional elongated error bars representing 1% and 99% percentiles. Colorcode: Red for African/African, blue for African/Non-African and orange for Non-African/Non-African pairs.

### Robustness to phasing and processing artifacts

MSMC2 (like MSMC) requires phased genomes for coalescence rate estimation, and we therefore rely on statistical phasing within the SGDP dataset, for which different strategies are possible. To compare the effect of selecting such phasing strategy, we generated phased datasets using eight different phasing strategies with three phasing algorithms (SHAPEIT[25], BEAGLE[26], EAGLE[27]). We included genotype calls from 12 individuals with previously published physically phased genomes[12] and then used those genomes to estimate the haplotype switch error rate. Among eight phasing strategies, SHAPEIT2[25], without the use of a reference panel, but including information from phase-informative reads[28], resulted in the lowest switch error rate per kb (and per heterozygous site; S8 Fig). Overall, switch error rates are higher in African populations, likely due to lower linkage disequilibrium, higher heterozygosity and relatively limited representation in the SGDP. To test how sensitive MSMC-IM is to different phasing strategies, we we tested four phasing strategies on the pair San/French. We find that the migration profile from MSMC-IM is very similar for different phasing strategies. In particular, we find that the very deep signal seen in population pairs involving San is consistent with different phasing strategies without a reference panel (S9 Fig). In a similar way, we tested the robustness of that signal with respect to choosing different filter levels (S9 Fig B) and with respect to removing CpG sites, which are known to have elevated mutation rates (S9 Fig C).

Given the superiority of the read-aware phasing strategy with SHAPEIT without a reference panel[28,29](S8 Fig), we used this method in all of our main analyses. However, even with this phasing strategy, the switch error rate is high in populations that are not well represented in the dataset. In case of indigenous Australians, the phasing quality is among the worst in the dataset (S8 Fig), arguably because the SGDP dataset contains only two Australian individuals (compared for example to 15 Papuans). To improve phasing in Australians specifically, we generated new high coverage genomic data for one of the two Australians in the SGDP dataset using a new library with longer read-pair insert sizes (see Methods). Using these additional reads reduced the switch error rate from 0.038/kb to 0.032/kb. (S8 Fig, blue isolated dot for Australian3). We ran MSMC2 on the long-insert Australian data, as well as the standard phased data, combined with one diploid genome from each of the other world-wide populations analysed in this study. The inferred migration profiles from MSMC-IM (S10 Fig) for Non-African population pairs involving the long-insert phased Australian genome do not seem to be affected by the phasing method (S10 Fig). The migration profile from pairs of Africans versus the long-insert phased Australian tend to be slightly younger, but also show deeper structure in Dinka/Australian, compared to the same pair using the *shapeit_pir* phasing method. Note that these migration rate densities exhibit more noise than the ones used in our main analysis (S4 Fig L), since they are based on only 1 individual per population, while the main analyses are based on two individuals per population. The main separation between Papuan and Australian remains at 15-35 kya, as shown in the migration profile from both phasing strategies, very close to the estimates from 8 haplotypes in the main analysis (S4 Fig L), and earlier than the previous estimates of 25-40kya[13].

Finally, to test internal consistency, we tested how well MSMC-IM was able to infer back its own model. We used the estimated migration rates and population sizes from five population pairs (Papuan/Australian, French/Han, Yoruba/French, French/Mbuti and French/San, 8 haplotypes per SGDP pair), and simulated genomic data under the inferred models for these population pairs. As shown in S11 Fig, the estimated migration patterns from the simulated and the real data are indeed very similar, including the deep signals seen in paris with San and Mbuti.

## Discussion

We have presented both a novel method MSMC-IM for investigating complex separation histories between populations, and an application of that method to human genomes, revealing new insights into the complex separations and deep ancestry in African populations. MSMC-IM extends MSMC2 by fitting an IM model to the estimated coalescence rates, which allows us to characterise the process of population separation via a continuous migration rate through time. In contrast to the established approach of using the relative cross coalescence rate directly from MSMC2, our new approach interprets coalescence rates more quantitatively. In a recent study a similar approach has been used to fit an IM model to PSMC estimates to estimate population split times and post-split migration rates in a more strictly parameterised model[30]. We found here that a continuous IM model without an explicit split time better fits the estimated coalescence rates from MSMC2, which are continuous themselves and thus lead to a more gradual concept of population separation. This absence of an explicit population split time distinguishes our approach from many previous models[5,8,9] and allows us to detect new signals of temporal population structure without specifying population phylogenies or admixture graphs from prior knowledge or via inference.

A showcase example for such new insights are the traces of extremely deep population structure seen in our analysis of African population pairs. The fact that San and Mbuti exhibit the deepest branches in the human population tree is itself not surprising given previous analyses[17,31–34], but the extraordinary time depth displayed in this analysis has to our knowledge not been reported before. This deep structure - albeit only making up 1% of ancestry - is far older than the oldest attested fossil records of anatomically modern humans, considering the East-African fossils of Omo Kibish and Herto 160-180 thousand years BP[34–36] and the skull from Jebel Irhoud recently re-dated to around 300 thousand years BP[37]. Any admixture from an archaic population that diverged from the main human lineage more than 600 thousand years go would produce such a signal. This is the case, for example, for the so-called “super-archaic” population that was inferred to have admixed into Denisovans[14] and was estimated to have diverged from the lineage leading to modern Humans, Neanderthals and Denisovans between 1.1 and 4 million years ago. Given this finding outside of Africa, it is perhaps not surprising that such deep archaic population structure existed also in Africa.

However, our signal of archaic population structure in Africa reveals more complexity than expected under the standard model of archaic introgression, in which two divergent populations admix with each other, creating a distinct pattern of deep ancestry in the genomes of the target population. Detecting such patterns in the genome would require a sufficient sequence divergence between non-introgressed and introgressed genomic segments (as measured by the S* statistic or extensions of it[15,16]). This is the case if the majority of ancestry between the two intermixing species has been isolated for hundreds of thousands of years, with a single split time. Such a scenario would be seen as a bimodal pattern in the migration profile reported by MSMC-IM, as shown in our simulations (Fig 2C). What we see, however, in the migration profiles between San and Mbuti with other African populations, is not a bimodal pattern, but a more continuous distribution. This would emerge under a model of repeated isolation and partial admixture of two or more archaic species or populations that exist in parallel for a long time. Under such a scenario, genomes are not a two-way mixture between introgressed and non-introgressed regions, but a mosaic of ancestry lines coalescing at a range of different split times. With no sharp boundary between introgressed and non-introgressed regions, methods such as S* fail to detect archaic ancestry, which may be the reason why the deep signals reported here have not been reported before for San and Mbuti, in contrast to Non-Africans and West-Africans[15,16].

While the continuous model in MSMC-IM adds significantly to previous approaches to estimating population separations, one drawback is that it is currently limited to only two populations at a time. While this limit is partially technical - MSMC2 cannot be scaled to arbitrary numbers of genomes - the more severe problem is a conceptual one. It is not obvious how to use the concept of continuous-time migration rates and non-sharp population separations to more than two populations. An important direction for future work is to achieve a generalisation of the continuous concept of population separation to multiple populations, which might help to better understand and quantify the processes that shaped human population diversity in the deep history of our species.

## Materials and Methods

### MSMC2

MSMC-IM is based on MSMC2 (first described and used in Ref. [13]) as a method to estimate pairwise coalescence rates from multiple genome sequences. The MSMC2 method is summarised in a self-contained way in S1 Text. MSMC2 is similar to MSMC [4], but instead of analysing multiple genomes simultaneously modelling the first coalescence event, it uses the pairwise model in sequence on all pairs of haplotypes to obtain a composite likelihood of the data given a demographic model. The demographic model itself (consisting of a piecewise constant coalescence rate) is then optimized via an Expectation-Maximization algorithm similarly to MSMC and PSMC [3]. For cross-population analyses, we use MSMC2 to obtain three independent coalescence rate estimates: two coalescence rates through time within each population, named λ_11_(*t*) and λ_22_(*t*), respectively, and one coalescence rate function for lineage pairs across the population boundary, named λ_12_(*t*) (S1 Text).

### MSMC-IM model

MSMC-IM then fits a two-island model with time-dependent population sizes *N*_1_(*t*) and *N*_2_(*t*), and a time-dependent continuous symmetric migration rate *m(t)* to the estimated coalescence rates, which essentially is a re-parameterization from the triple of functions {λ_11_(*t*), λ_12_(*t*), λ_22_(*t*)} to a new triple of functions {*N_1_(t), N_2_(t), m(t)*} (S1 Text). To fit the island-model to the coalescence rates, we first use the coalescence rates to compute a probability density for times to the most recent common ancestor (tMRCA), as illustrated here for rate λ_11_(*t*):

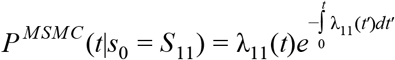

Here, *S*_11_ denotes the starting state where both lineages are present in population 1. We then use an approach by Hobolth et al 2011 [38] to compute this density for the three starting states *s*_0_ = {*S*_11_,*S*_12_, *S*_22_} under an IM model, denoted *P^IM^*(*t*|*s*_0_), using exponentiation of the rate matrix of the underlying IM-Markov process that governs the state of uncoalesced and coalesced lineages in two populations connected by a time-dependent migration rate (see S1 Text). The fitting process of the IM model to the density computed from MSMC2 is done by minimizing the Chi-square statistics:

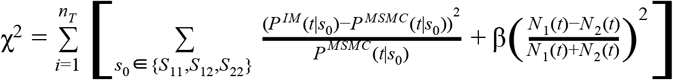

Where the second term is a regularization term to avoid overfitting, pushing the two population sizes *N_1_(t)* and *N_2_(t)* close to each other. The strength of this regularization can be controlled via a user-defined parameter in our program. For the three simulation scenarios and all pairs of real data expect pairs including Han, we used a regularisation value of 10^−6^. Regularization is necessary because the reparameterization introduced by MSMC-IM overspecifies the model at times when the two populations are fully merged. For that same reason, we plot estimated migration rates in all figures only up to a value of M(t) = 0.999, since migration rate estimates beyond that point are essentially arbitrary, as lineages have already been fully randomized between the two populations. We also restrict the estimated population sizes to 10,000,000 in practice.

We implemented the MSMC-IM model as a python command line utility that takes the MSMC output files as input. The program is available at: https://github.com/wangke16/MSMC-IM

### Simulations

We used msprime[39] for all simulations in this paper. In the three series of simulation scenarios mentioned above, we simulated four diploid genomes composed of 22 chromosomes each of length 100Mbp from two populations, assuming a constant population size 20,000 for every population. The recombination rate we used here is 10^−8^ per generation per bp, and the mutation rate is 1.25 × 10^−8^.

In the zig-zag simulation (S1 Fig), we simulated a series of exponential population growths and declines for two, four and eight haplotypes, each changing between 3,000 and 30,000 in exponentially increasing time intervals, with the same simulation parameters as specified in Ref. [4] and Ref.[3] to ensure comparability with these previous publications. In particular, this simulation involved a lower recombination rate (0.3 × 10^−8^) than the main simulations, justified in [4] as the inferred recombination rate from real data using PSMC’. The reason for it being lower than the true recombination rate (close to 10^−8^, as used in the main simulations above), is that MSMC (and MSMC2) infers an “effective recombination rate”, which is a non-trivial average over the variable recombination landscape across the human genome.

We also conducted a number of simulations based on MSMC-IM inference from real data (S11 Fig). We took the estimates on migration rates and population sizes from MSMC-IM (S2 Table) for five pairs of worldwide populations (San/French, Mbuti/French, Yoruba/French, French/Han, Papuan/Australian), as the input parameters in our simulation, and simulated 2.2Gb genomes on 8 haplotypes for each case.The recombination rate we used here is 10^−8^ per generation per bp, and the mutation rate is 1.25 × 10^−8^.

### World-wide genomic Data

For the results shown in Figs 3-6, we used 30 high coverage genomes from 15 cross-continental modern populations in the SGDP dataset[12], with two diploid genomes from each population for running MSMC2 and MSMC-IM (S1 Table). We ran pairwise analyses for 13 populations (excluding Quechua and Mixe) and pairwise comparisons within three native American populations (81 population pairs in total). We downloaded the cteam-lite dataset of from the website: http://reichdata.hms.harvard.edu/pub/datasets/sgdp/, in the hetfa-format where all sites are represented by an IUPAC encoding representing diploid genotypes, along with individual masks recording the quality of the genotype calls. We converted the hetfa mask files (.ccompmask.fa.rz) to zipped bed format though two steps: first we uncompressed the hetfa mask files using “*htsbox razip -d -c*” (https://github.com/lh3/htsbox), and then converted the uncompressed mask files (.ccompmask.fa) to zipped bed format by an in-house python script adapted from the *makeMappabilityMask.py* script in *msmc-tools* (www.github.com/stschiff/msmc-tools). The cteam-lite masks encode quality using an integer-range from 0 to 9 (reflecting increasing stringency) and “N” to represent missing data. For our analysis, we included all sites that were non-missing, i.e. have a minimum quality level of 0.

Following the processing introduced in PSMC [3] and MSMC/MSMC2 [4], beyond the individual masks we also use a universal mask to reflect overall mappability and SNP calling properties along the human genome. We used the universal masks defined in Supplementary Info 4 from Ref. [12] (and available for download at https://github.com/wangke16/MSMC-IM/masks) as additional negative masks denoting genomic regions to be filtered out.

Beside the genome-wide mask files for each individual, we obtained variant data as made available on the SGDP project website (https://sharehost.hms.harvard.edu/genetics/reich_lab/sgdp/phased_data/). Due to the specifics of how that dataset was generated, only segregating sites at positions where the Chimpanzee reference genome has non-missing data are included. To balance this missigness based on the Chimpanzee reference genome for MSMC, we included an additional mask in our preprocessing, which reflected non-missing regions in the Chimpanzee reference sequence. For others to reproduce our analysis, we provide this chimp mask on the MSMC-IM github repository (https://github.com/wangke16/MSMC-IM). We conducted several runs with removed CpG sites. For this, we generated a mask including all positions of Cytosines and Guanines in CpG dinucleotides, Thymines in TpG dinucleotides, and Adenosines in CpA dinucleotides in the human reference genome hg19, and used those positions as negative mask when preparing the MSMC input files. This mask can be found in the same github repository as above.

We phased the data using SHAPEIT2 (v837) [25], Beagle4.0 (r1399) [26] and EAGLE2 (version 2.3) [27]. We first phased the data using each algorithm both with and without a reference panel. When using a reference panel, all three methods are only able to phase sites that are represented in the reference panel. Therefore, we removed sites not in the reference panel, phased, adding the removed sites back as unphased, and then ran a second round of phasing using Beagle4.0 and the “usephase=true” option, which allows us to phase the unphased sites in data that is already partially phased. Finally, we also phased using SHAPEIT2 without a reference panel, but using the read-aware phasing stratergy[28]. This uses the fact that two SNPs found on the same (paired) read must be in phase. The switch error of each of these phasing strategies, evaluated by comparison with the experimentally phased data generated for the same samples[12] is shown in S8 Fig.

Finally, we generated a long-insert library from one of the two Australian DNA samples analysed in SGDP [12], with a median insert size of 3.3kbp. This data is available at the European Nucleotide Archive under accession number ERX1790596 (https://www.ebi.ac.uk/ena/data/view/ERX1790596). We used this data to improve the phasing quality for this Australian individual. As shown in S6 Fig, this strategy indeed reduced the switch error rate for this Australian individual from 0.036/kb to 0.032/kb.

### Running MSMC-IM

Unlike MSMC, which reports these three rates in a single analysis step, in MSMC2 we run the three estimations for λ_11_(*t*), λ_12_(*t*) and λ_22_(*t*) independently from each other, using a different selection of haplotype pairs in each case. We base most of our analyses on 4 diploid individuals (unless indicated otherwise), for which we prepared joint input files for each chromosome, consisting of 8 haplotypes each. We then chose the pairs to be analysed using the “-I” option in MSMC2. For coalescence rate λ_11_(*t*), we used “-I 0,1,2,3”, which instructs MSMC2 to iterate through all six possible haplotype pairs among the four haplotypes from the first population. Likewise, to estimate λ_22_(*t*), we used “-I 4,5,6,7”. Finally, to obtain estimates of the coalescence rates across populations, λ_12_(*t*), we used “-I 0-4,0-5,0-6,0-7,1-4,1-5,1-6,1-7,2-4,2-5,2-6,2-7,3-4,3-5,3-6,3-7”, iterating through all sixteen possible haplotype pairings between the four haplotypes in each population.MSMC-IM requires a single input file containing all three coalescence rate estimates, similar to the output generated by the original MSMC program. A script *combineCrossCoal.py*is provided on the msmc-tools github repository (http://www.github.com/stschiff/msmc-tools), to generate the combined output file from the three output files of the three MSMC2 runs for a pair of populations

With the combined MSMC2 output as input, we run MSMC-IM model by “***MSMC_IM.py pair.combined.msmc2.txt***”. Also the time pattern needs to be specified, which is by default *1*2+25*1+1*2*+*1*3* as the default in MSMC2. In the output, MSMC-IM will rescale the scaled time in MSMC2 output by mutation rate 1.25e-8 into real time in generations, and report symmetric migration rates and M(t) in each time segment.

### Robustness of Phasing strategies

We tested the robustness of our results by applying different phasing strategies and mask filtering levels to a single pair of San/French in the SGDP dataset. The phasing strategy was varied between *beagle, shapeit, shapeit_ref_all* to *shapeit_pir.* Here, *beagle* and *shapeit* denote phasing with no reference panel, *shapeit_ref_all* denotes phasing with a reference panel (with sites not in the reference panel phased with Beagle) and shapeit_pir denotes no reference panel but including phase-informative reads.. The stringency of the filtering mask was varied between filter levels 0, 1, 3, 5. Beyond that, we tested our approach on 12 populations (24 genomes) from another dataset [14], which consists of different genomes. This dataset was processed independently using the pipeline in the msmc-tools github repository (http://www.github.com/stschiff/msmc-tools) i.e. SNPs and masks generated using samtools and *bamCaller.py*, with statistical phasing by *SHAPEIT2* with the 1000 Genomes reference panel, leaving sites not present in the reference panel as unphased.

## Acknowledgements

We thank Erich Jaeger and Aparna Natarajan at Illumina for generating the Australian Nextera Mate Pair library and providing sequencing data. SS and KW acknowledge support by the Max Planck Society. IM was supported by a Sloan Research Fellowship and a New Investigator Research Grant from the Charles E. Kaufman fund of The Pittsburgh Foundation.

## Author Contributions

KW and SS developed the model. KW implemented and tested the model. KW and IM processed data. KW and SS analysed the data. JOC and IM supervised data-generation and bioinformatic processing of the long-insert Australian genomic data. SS supervised the study. KW and SS wrote the paper, with input from all authors.

## References

1. McVean GAT, Cardin NJ. Approximating the coalescent with recombination. Philos Trans R Soc Lond B Biol Sci. 2005;360: 1387–1393.

2. Marjoram P, Wall JD. Fast “coalescent” simulation. BMC Genet. 2006;7: 16.

3. Li H, Durbin R. Inference of human population history from individual whole-genome sequences. Nature. 2011;475: 493–496.

4. Schiffels S, Durbin R. Inferring human population size and separation history from multiple genome sequences. Nat Genet. Nature Publishing Group; 2014;46: 919–925.

5. Steinrücken M, Kamm JA, Song YS. Inference of complex population histories using whole-genome sequences from multiple populations [Internet]. Cold Spring Harbor Labs Journals; 2015 Sep. Available: http://biorxiv.org/lookup/doi/10.1101/026591

6. Sheehan S, Harris K, Song YS. Estimating variable effective population sizes from multiple genomes: a sequentially markov conditional sampling distribution approach. 2013;194: 647–662.

7. Terhorst J, Kamm JA, Song YS. Robust and scalable inference of population history from hundreds of unphased whole genomes. Nat Genet. Nature Publishing Group; 2017;49: 303–309.

8. Kamm JA, Terhorst J, Song YS. Efficient computation of the joint sample frequency spectra for multiple populations. J Comput Graph Stat. 2017;26: 182–194.

9. Kamm JA, Terhorst J, Durbin R, Song YS. Efficiently inferring the demographic history of many populations with allele count data. doi:10.1101/287268

10. Excoffier L, Foll M. fastsimcoal: a continuous-time coalescent simulator of genomic diversity under arbitrarily complex evolutionary scenarios. Bioinformatics. 2011;27: 1332–1334.

11. Schiffels S, Haak W, Paajanen P, Llamas B, Popescu E, Loe L, et al. Iron Age and Anglo-Saxon genomes from East England reveal British migration history. Nat Commun. 2016;7: 10408.

12. Mallick S, Li H, Lipson M, Mathieson I, Gymrek M, Racimo F, et al. The Simons Genome Diversity Project: 300 genomes from 142 diverse populations. Nature. 2016;538: 201–206.

13. Malaspinas A-S, Westaway MC, Muller C, Sousa VC, Lao O, Alves I, et al. A genomic history of Aboriginal Australia. Nature. 2016;538: 207–214.

14. Prüfer K, Racimo F, Patterson N, Jay F, Sankararaman S, Sawyer S, et al. The complete genome sequence of a Neanderthal from the Altai Mountains. Nature. 2014;505: 43–49.

15. Plagnol V, Wall JD. Possible ancestral structure in human populations. PLoS Genet. 2006;2: e105.

16. Durvasula A, Sankararaman S. Recovering signals of ghost archaic admixture in the genomes of present-day Africans [Internet]. bioRxiv. 2018. p. 285734. doi:10.1101/285734

17. Pickrell JK, Patterson N, Barbieri C, Berthold F, Gerlach L, Güldemann T, et al. The genetic prehistory of southern Africa. Nat Commun. 2012;3: 1143.

18. Skoglund P, Thompson JC, Prendergast ME, Mittnik A, Sirak K, Hajdinjak M, et al. Reconstructing Prehistoric African Population Structure. Cell. 2017;171: 59–71.e21.

19. Sankararaman S, Mallick S, Dannemann M, Prüfer K, Kelso J, Pääbo S, et al. The genomic landscape of Neanderthal ancestry in present-day humans. Nature. 2014;507: 354–357.

20. Sankararaman S, Mallick S, Patterson N, Reich D. The Combined Landscape of Denisovan and Neanderthal Ancestry in Present-Day Humans. Curr Biol. 2016;26: 1241–1247.

21. Browning SR, Browning BL, Zhou Y, Tucci S, Akey JM. Analysis of Human Sequence Data Reveals Two Pulses of Archaic Denisovan Admixture. Cell. 2018;173: 53–61.e9.

22. Pagani L, Lawson DJ, Jagoda E, Mörseburg A, Eriksson A, Mitt M, et al. Genomic analyses inform on migration events during the peopling of Eurasia. Nature. 2016;538: 238–242.

23. Meyer M, Kircher M, Gansauge M-T, Li H, Racimo F, Mallick S, et al. A high-coverage genome sequence from an archaic Denisovan individual. Science. 2012;338: 222–226.

24. Raghavan M, Skoglund P, Graf KE, Metspalu M, Albrechtsen A, Moltke I, et al. Upper Palaeolithic Siberian genome reveals dual ancestry of Native Americans. Nature. 2014;505: 87–91.

25. Delaneau O, Zagury J-F, Marchini J. Improved whole-chromosome phasing for disease and population genetic studies. Nat Methods. 2013;10: 5–6.

26. Browning SR, Browning BL. Rapid and accurate haplotype phasing and missing-data inference for whole-genome association studies by use of localized haplotype clustering. Am J Hum Genet. 2007;81: 1084–1097.

27. Loh P-R, Danecek P, Palamara PF, Fuchsberger C, A Reshef Y, K Finucane H, et al. Reference-based phasing using the Haplotype Reference Consortium panel. Nat Genet. 2016;48: 1443–1448.

28. Delaneau O, Howie B, Cox AJ, Zagury J-F, Marchini J. Haplotype estimation using sequencing reads. Am J Hum Genet. 2013;93: 687–696.

29. Choi Y, Chan AP, Kirkness E, Telenti A, Schork NJ. Comparison of phasing strategies for whole human genomes. PLoS Genet. 2018;14: e1007308.

30. Song S, Sliwerska E, Emery S, Kidd JM. Modeling Human Population Separation History Using Physically Phased Genomes. Genetics. 2017;205: 385–395.

31. Tishkoff SA, Gonder MK, Henn BM, Mortensen H, Knight A, Gignoux C, et al. History of click-speaking populations of Africa inferred from mtDNA and Y chromosome genetic variation. Mol Biol Evol. 2007;24: 2180–2195.

32. Knight A, Underhill PA, Mortensen HM, Zhivotovsky LA, Lin AA, Henn BM, et al. African Y chromosome and mtDNA divergence provides insight into the history of click languages. Curr Biol. 2003;13: 464–473.

33. Schlebusch CM, Skoglund P, Sjödin P, Gattepaille LM, Hernandez D, Jay F, et al. Genomic variation in seven Khoe-San groups reveals adaptation and complex African history. Science. 2012;338: 374–379.

34. Schlebusch CM, Jakobsson M. Tales of Human Migration, Admixture, and Selection in Africa. Annu Rev Genomics Hum Genet. 2018; doi:10.1146/annurev-genom-083117-021759

35. McDougall I, Brown FH, Fleagle JG. Stratigraphic placement and age of modern humans from Kibish, Ethiopia. Nature. 2005;433: 733–736.

36. White TD, Asfaw B, DeGusta D, Gilbert H, Richards GD, Suwa G, et al. Pleistocene Homo sapiens from Middle Awash, Ethiopia. Nature. 2003;423: 742–747.

37. Richter D, Grün R, Joannes-Boyau R, Steele TE, Amani F, Rué M, et al. The age of the hominin fossils from Jebel Irhoud, Morocco, and the origins of the Middle Stone Age. Nature. 2017;546: 293–296.

38. Hobolth A, Andersen LN, Mailund T. On computing the coalescence time density in an isolation-with-migration model with few samples. Genetics. 2011;187: 1241–1243.

39. Kelleher J, Etheridge AM, McVean G. Efficient Coalescent Simulation and Genealogical Analysis for Large Sample Sizes. PLoS Comput Biol. 2016;12: e1004842.

