## Supplementary Figures and Tables for "Tracking human population structure through time from whole genome sequences"

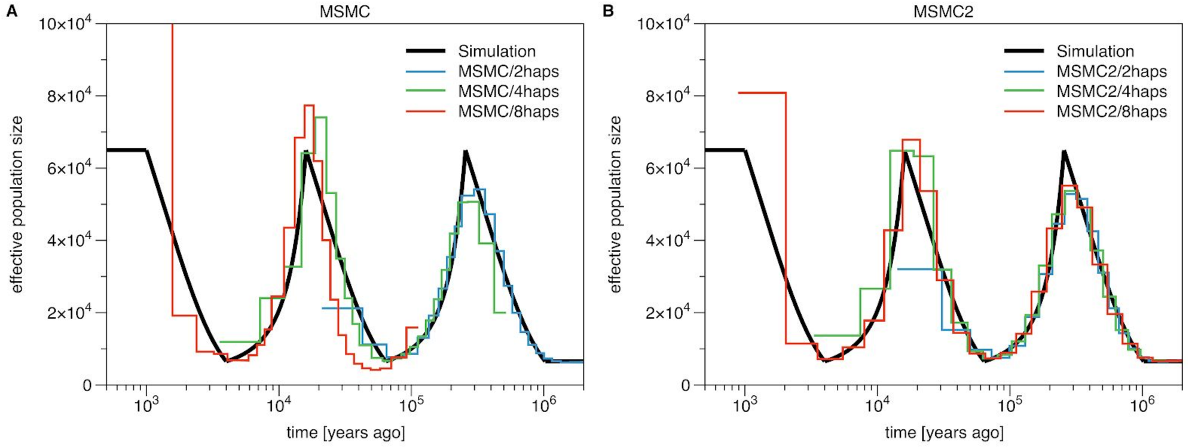


**S1 Figure. MSMC and MSMC2 population size estimates from simulated data.** To test the resolution of MSMC (A) and MSMC2 (B) applied to two, four and eight haplotypes, we simulated a series of exponential population growths and declines, each changing the population size by a factor ten. The true population size is shown as dark solid line. Compared to MSMC, MSMC2 recovers the population size well, and the resolution in recent times increases with the number of haplotypes. With two haplotypes, MSMC2 infers the population history from 10,000 years ago to 3 million years, whereas, with four haplotypes and eight haplotypes the resolution in recent times is extended to 3,000 and 1,000 years ago respectively.

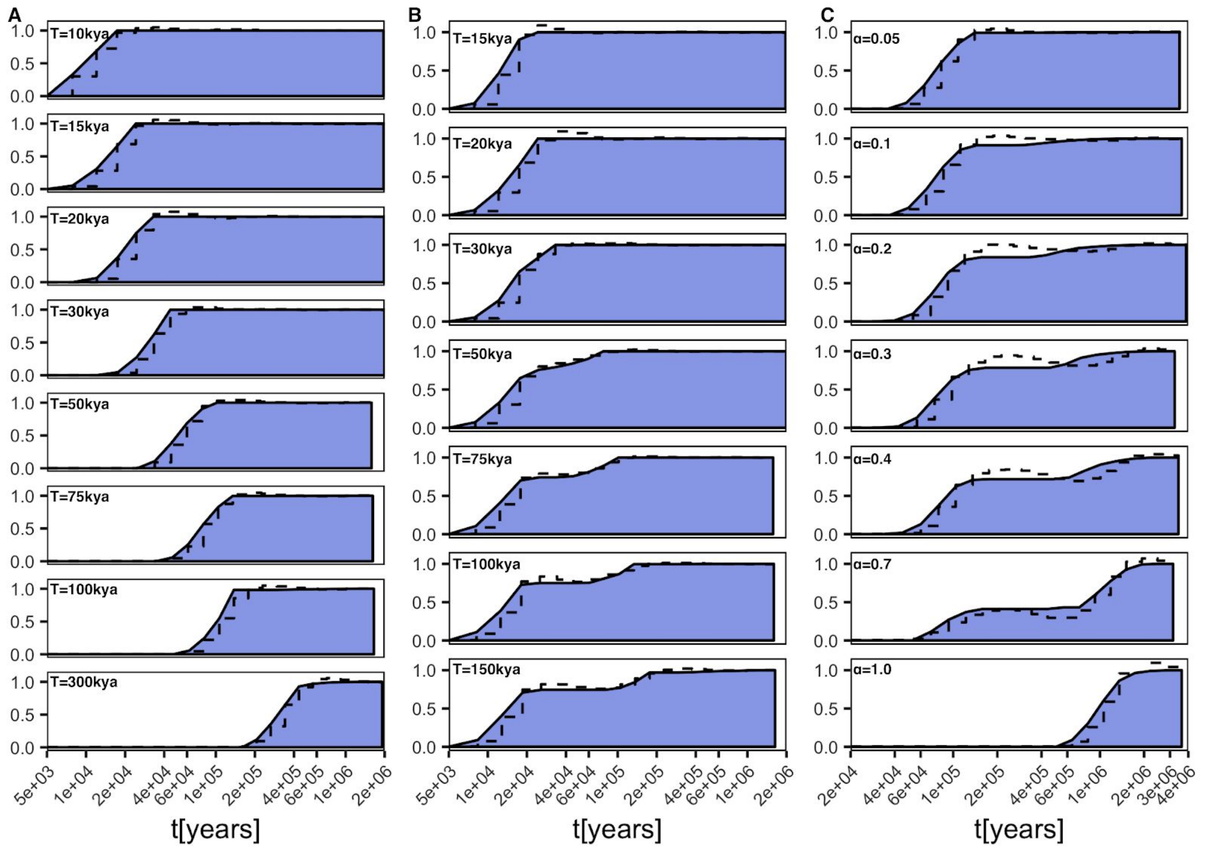


**S2 Figure. Cumulative migration densities from three simulation scenarios.** This figure shows the same results as Figure 2, but showing M(t) instead of m(t). The scenarios are (A) the *Clean-split-scenario*. (B) the *Split-with-migration*-scenario, and (C) the *Split-with-archaic-admixture*-scenario.

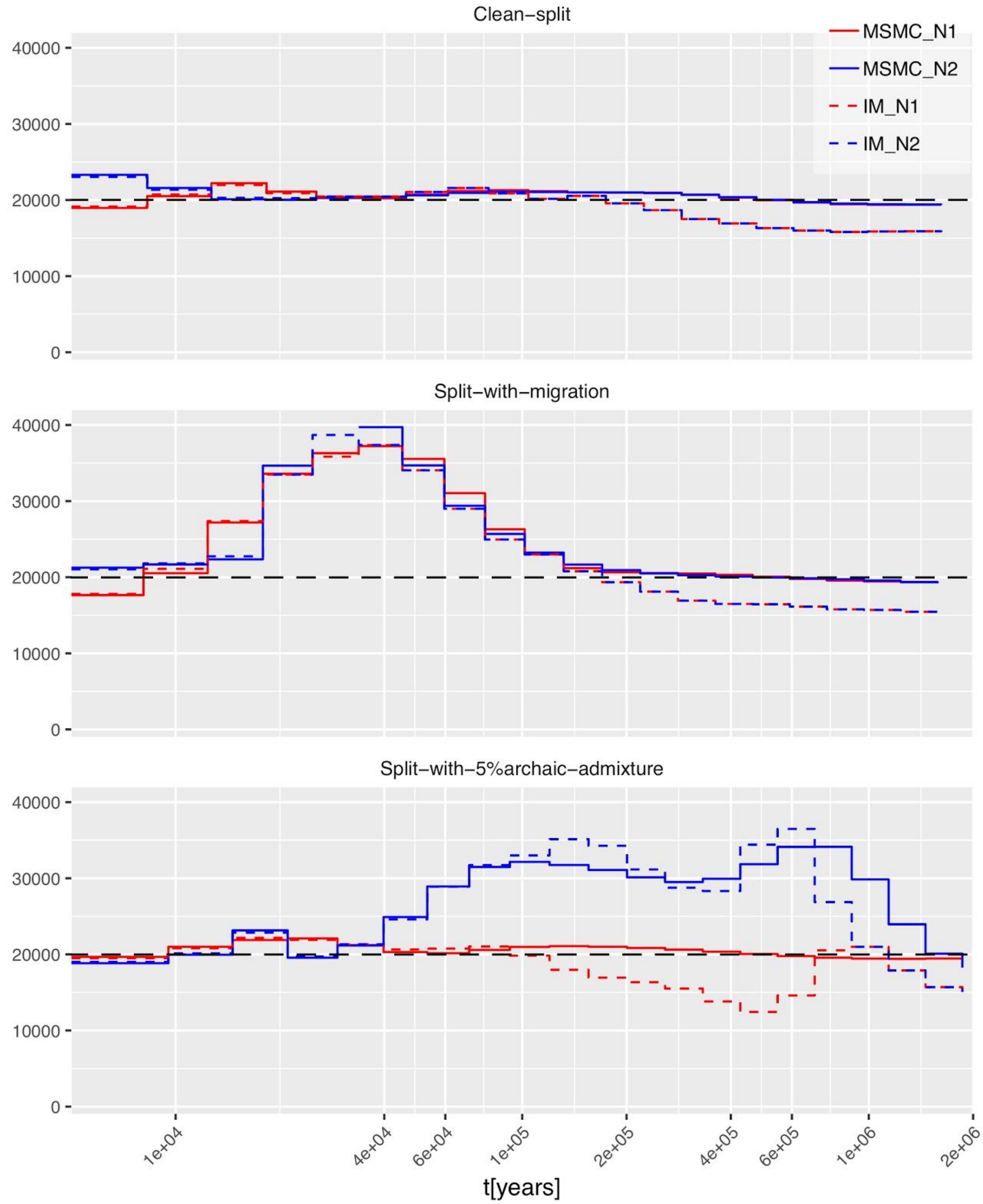


**S3 Figure. Population size estimates from MSMC2 compared to MSMC-IM:** we simulated $N_1(t)$ and $N_2(t)$ as constant 20,000 (shown as black dashed lines) in three different simulation scenarios. The split time T is at 75kya in all three cases, and all other parameters are the same as in Figure 2. As shown, the MSMC-IM estimates for $N_1(t)$ and $N_2(t)$ are close to the inverse coalescence rates, without much effect from the estimated migration rates.

**S4 Figure. Pairwise migration profiles for 13 worldwide populations:** San (A), Mbuti (B), Mandenka (C), Dinka (D), Yoruba (E), Mende (F), French (G), Sardinian (H), Han (I), Dai (J), Papuan (K), Australian (L), Karitiana (M). *See separate joint PDF file.*

**S5 Figure. Migration profile of an independent dataset.** Here we have analysed 12 worldwide populations from Prüfer et al (2014) with independent data processing as described in Methods: San (A), Mbuti (B), Mandenka (C), Dinka (D), Yoruba (E), French (F), Sardinian (G), Han (H), Dai (I), Papuan (J), Australian (K), Karitiana (L). *See separate joint PDF file*.

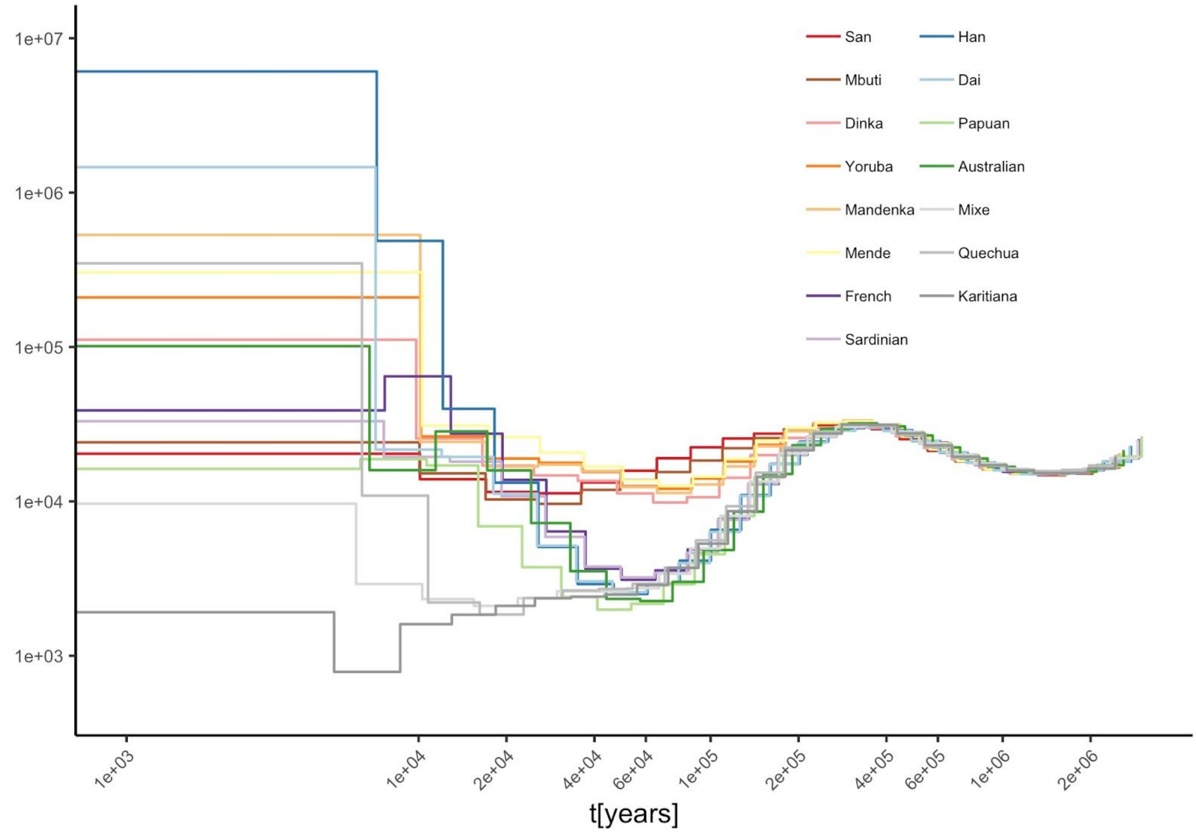


**S6 Figure. Estimated population sizes from MSMC2 for 15 worldwide populations.**

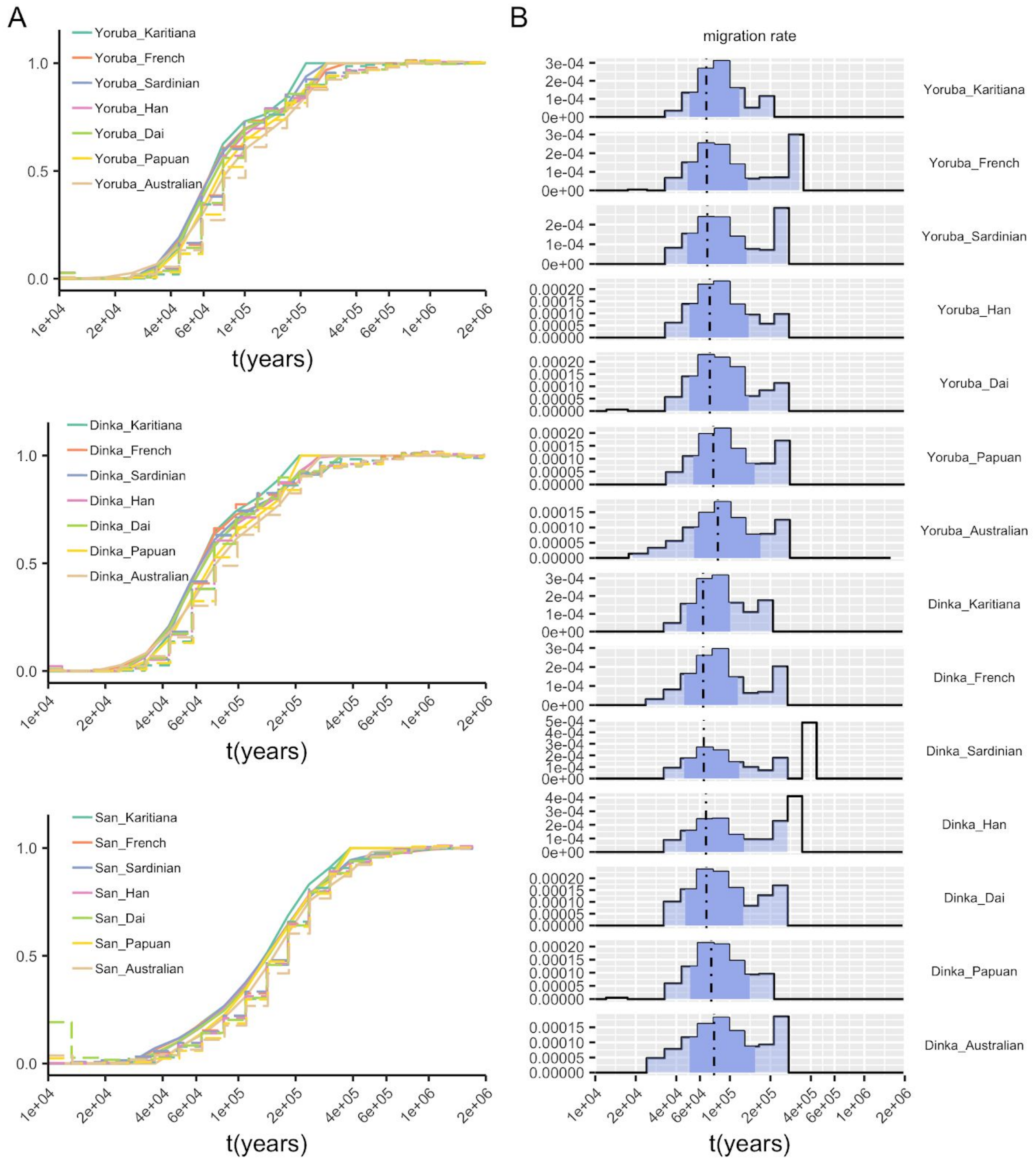


**S7 Figure. Testing for multiple out-of-Africa scenarios.** Here we show analyses on the divergence of Papuans and Australians from Africa vs. other Non-African populations from Africa. We show the Relative Cross Coalescence Rate (rCCR, dashed) and the cumulative migration density, M(t) (solid) (A), and the migration density m(t) (B) for pairs of populations of Yoruba, Dinka and San with one non-African population as indicated.

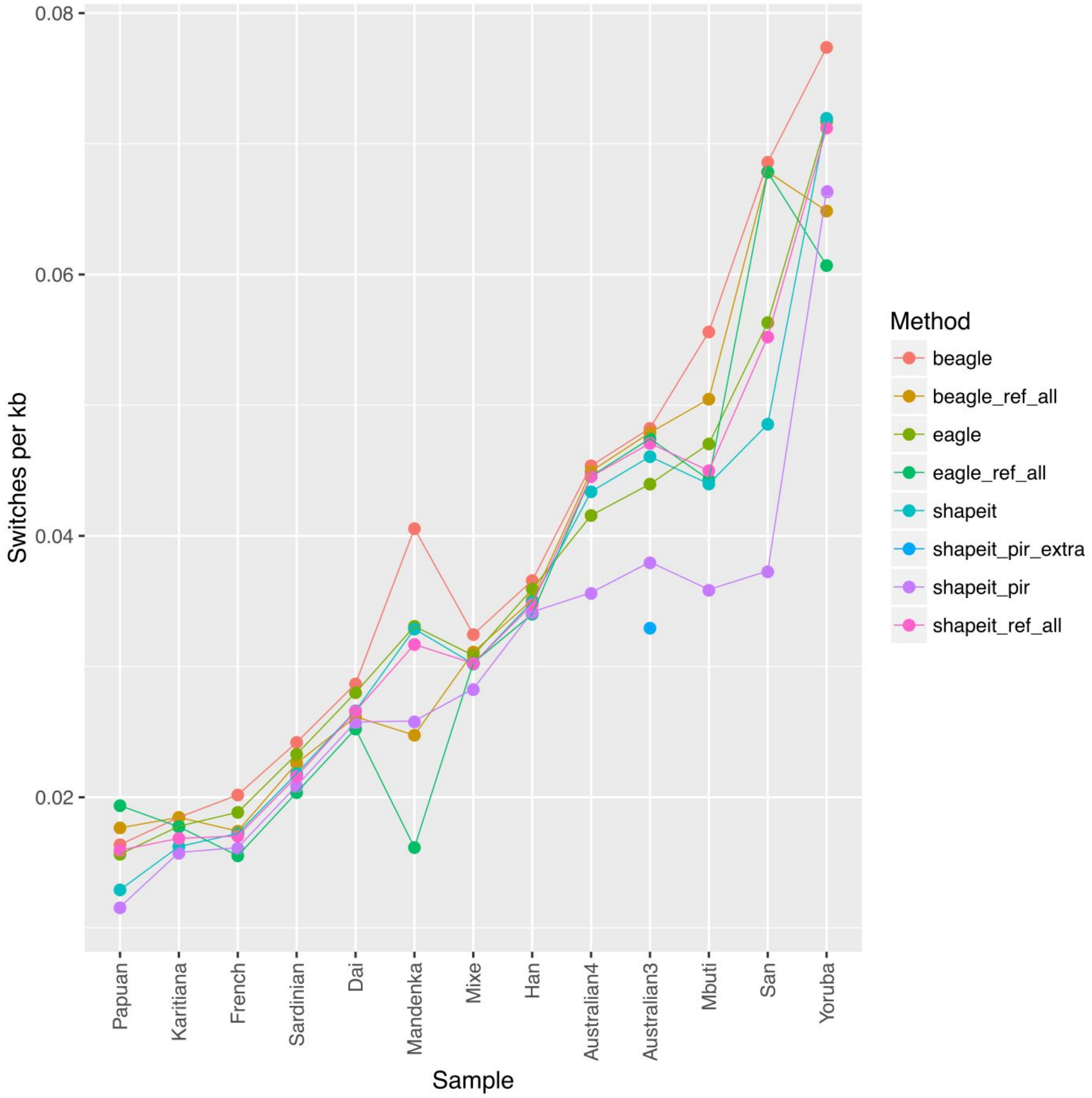


**S8 Figure. Switch error rates from eight phasing strategies**. *beagle* and *beagel_ref_all* denote BEAGLE phasing without and with reference panel. *eagle* and *eagle_ref_all* represent EAGLE phasing without and with reference panel. *shapeit* and *shapeit_ref_all* represent SHAPEIT phasing without and with reference panel. *shapeit_pir* represents SHAPEIT phasing with phase-informative reads. *shapeit_pir_extra* represents SHAPEIT phasing with long-insert-size reads as phase informative reads, which was applied to B-Australian-3 only. See Methods for details.

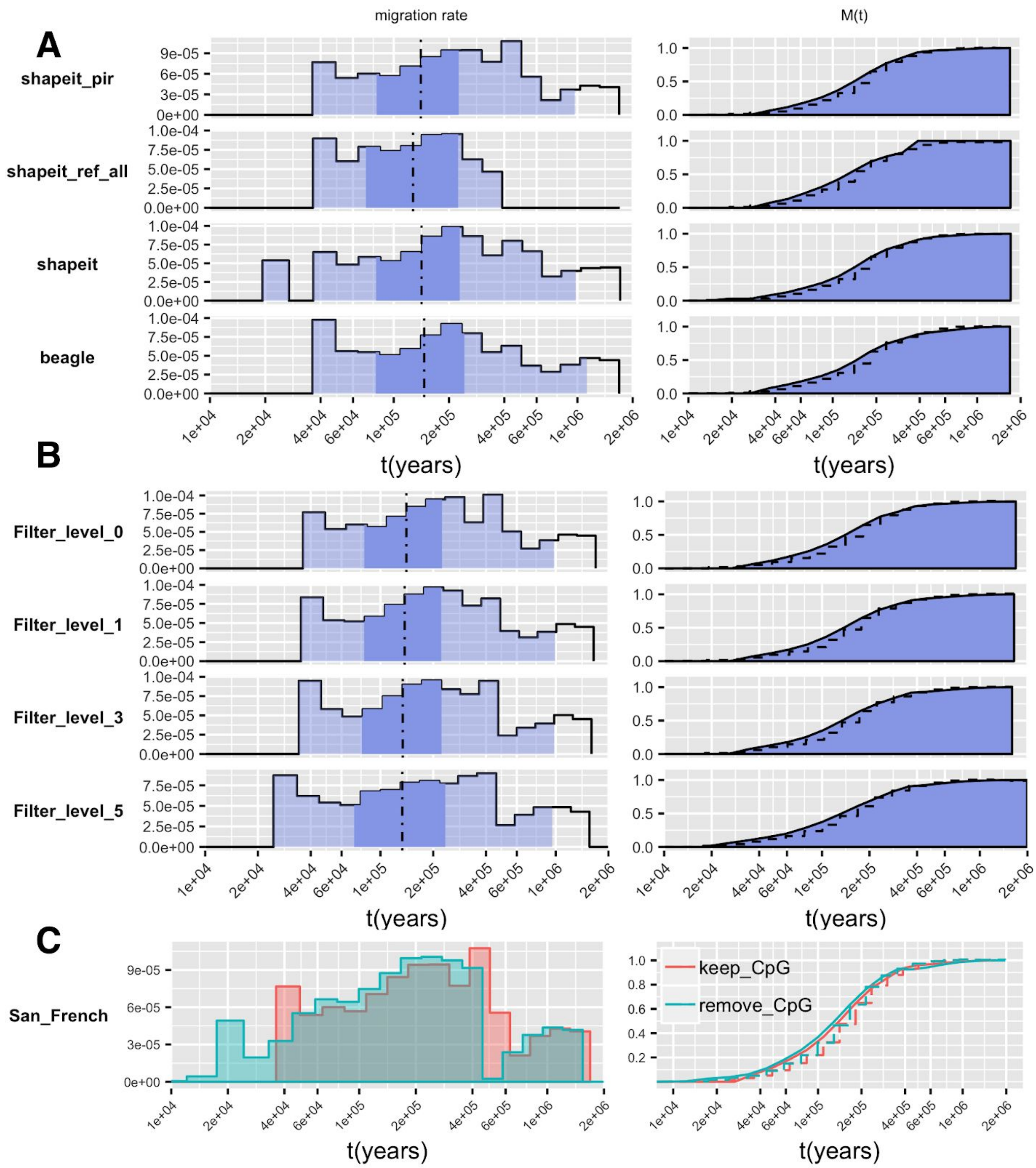


**S9 Figure. Impact of phasing and processing artifacts.** Using results from the pair San/French as an example, we show (A) the impact of the phasing strategy, (B) the impact of the filtering level for generating individual masks, and (C) the impact of removing CpG sites.

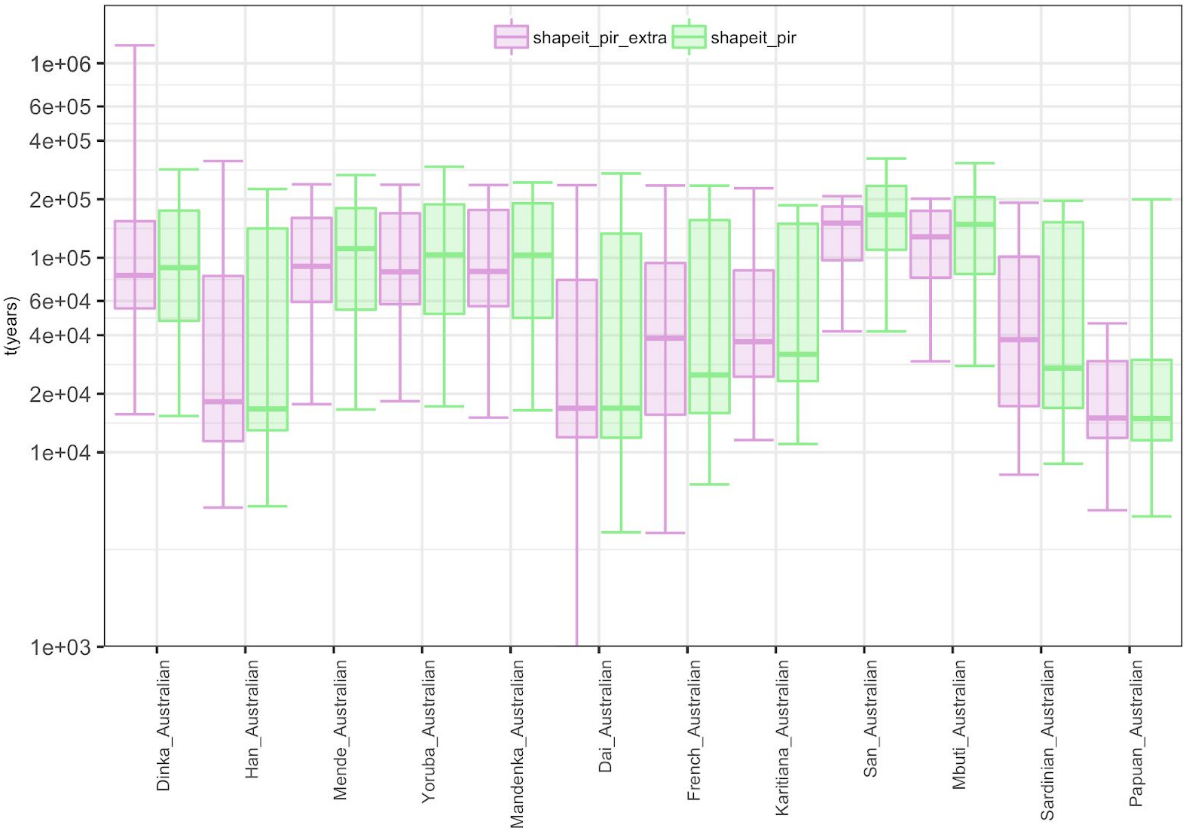


**S10 Figure. Impact of long-insert phasing on Australian-involved population separation inferences**. M(t) in quantiles is summarised here between a single Australian and a single individual from worldwide populations**.** Boxes show the 25% to 75% quantiles of M(t), with bi-directional elongated error bars representing 1% and 99% percentiles. Purple color represents the data phased using long-insert reads. Green color represents the standard phased dataset.

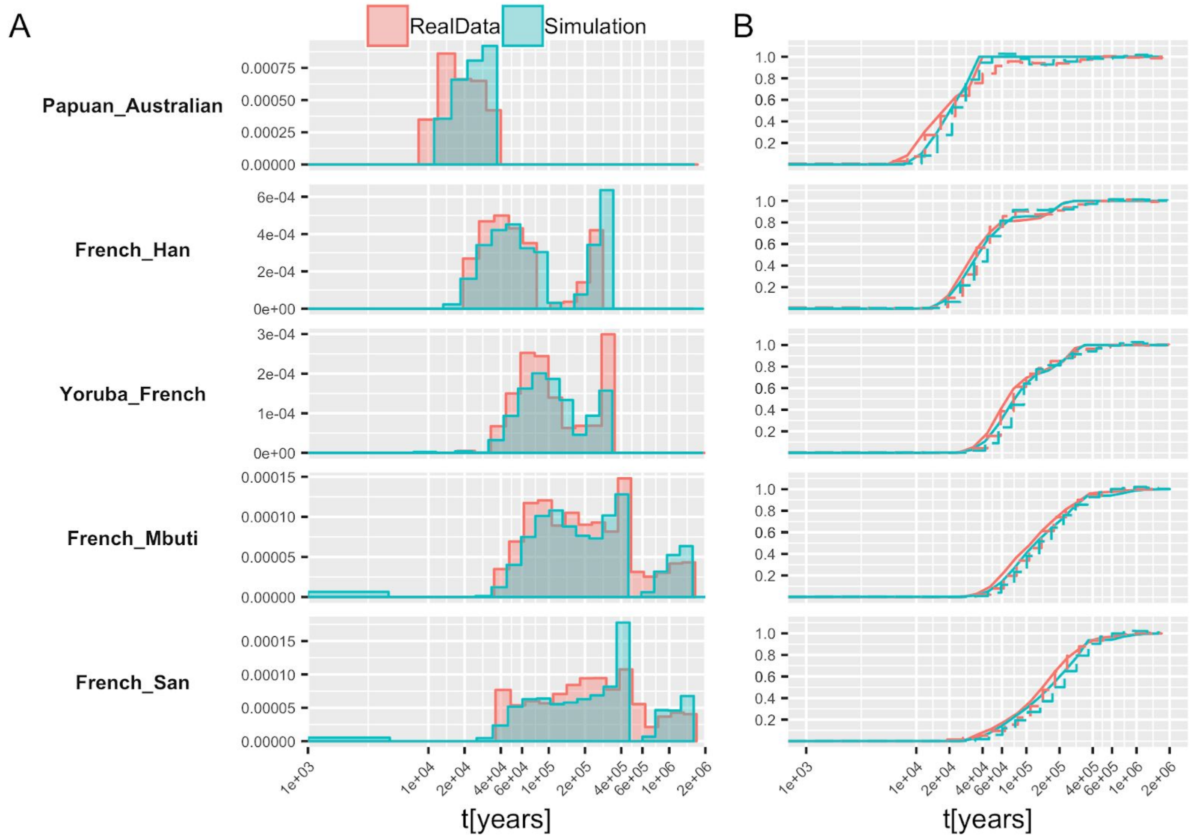


**S11 Figure. Migration profile on simulated pseudo-SGDP genomes.** Red color shows the estimates we got from SGDP dataset for pairs shown on the left, which is the true parameter used in the simulation. Green color shows the estimates from simulated genomes given the true parameter shown in red. (A) Migration rates. (B) M(t) and rCCR changes along time.

**S1 Table.** Analysed samples and population labels from the SGDP dataset.

| Sample ID | Population Label | Continent |
|---|---|---|
| S_Yoruba-1 | Yoruba | Africa |
| S_Yoruba-2 | Yoruba | Africa |
| S_Dinka-1 | Dinka | Africa |
| S_Dinka-2 | Dinka | Africa |
| S_Mbuti-1 | Mbuti | Africa |
| S_Mbuti-2 | Mbuti | Africa |
| S_Mandenka-1 | Mandenka | Africa |
| S_Mandenka-2 | Mandenka | Africa |
| S_Mende-1 | Mende | Africa |
| S_Mende-2 | Mende | Africa |
| S_Khomani_San-1 | San | Africa |
| S_Khomani_San-1 | San | Africa |

| | | |
|---|---|---|
| S_Sardinian-1 | Sardinian | Europe |
| S_Sardinian-2 | Sardinian | Europe |
| S_French-1 | French | Europe |
| S_French-2 | French | Europe |
| S_Han-1 | Han | East Asia |
| S_Han-2 | Han | East Asia |
| S_Dai-1 | Dai | East Asia |
| S_Dai-2 | Dai | East Asia |
| S_Papuan-1 | Papuan | New Guinea |
| S_Papuan-2 | Papuan | New Guinea |
| B_Australian-3 | Australian | Australia |
| B_Australian-4 | Australian | Australia |
| S_Karitiana-1 | Karitiana | South America |
| S_Karitiana-2 | Karitiana | South America |
| S_Quechua-1 | Quechua | South America |
| S_Quechua-2 | Quechua | South America |
| S_Mixe-2 | Mixe | South America |
| S_Mixe-3 | Mixe | South America |

**S2 Table**. Table of MSMC2 results and MSMC-IM estimates for all pairs of SGDP populations analysed, *see separate Excel file.*

**S1 Text**: Derivation of MSMC2 and MSMC-IM theory, *see separate PDF file.*
