## Supplementary Fig5 for "Tracking human population structure through time from whole genome sequences"

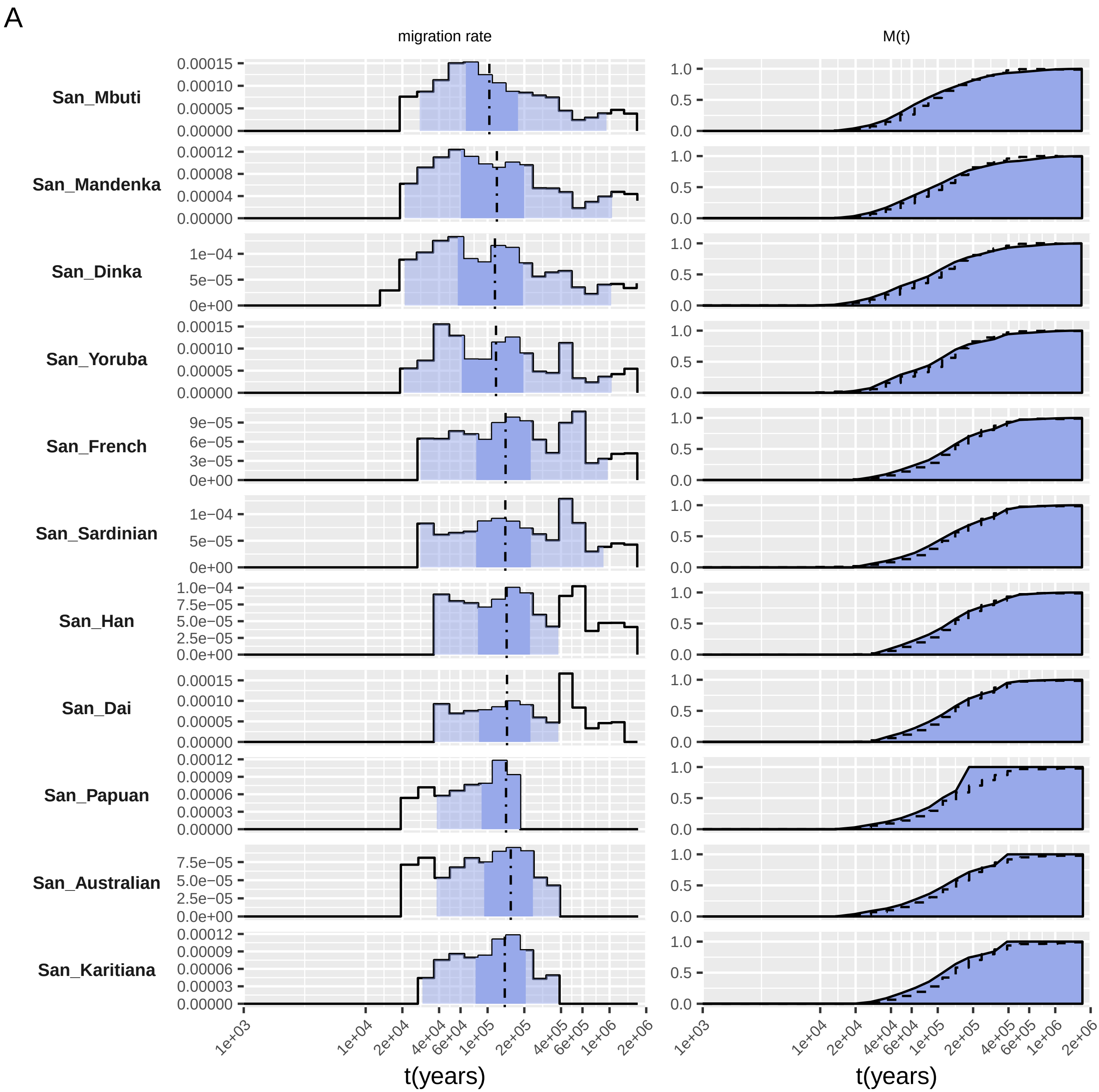

A
migration rate
M(t)
San_Mbuti
San_Mandenka
San_Dinka
San_Yoruba
San_French
San_Sardinian
San_Han
San_Dai
San_Papuan
San_Australian
San_Karitiana
1e+03
1e+04
2e+04
4e+04
6e+04
1e+05
2e+05
4e+05
6e+05
1e+06
2e+06
t(years)
t(years)

B

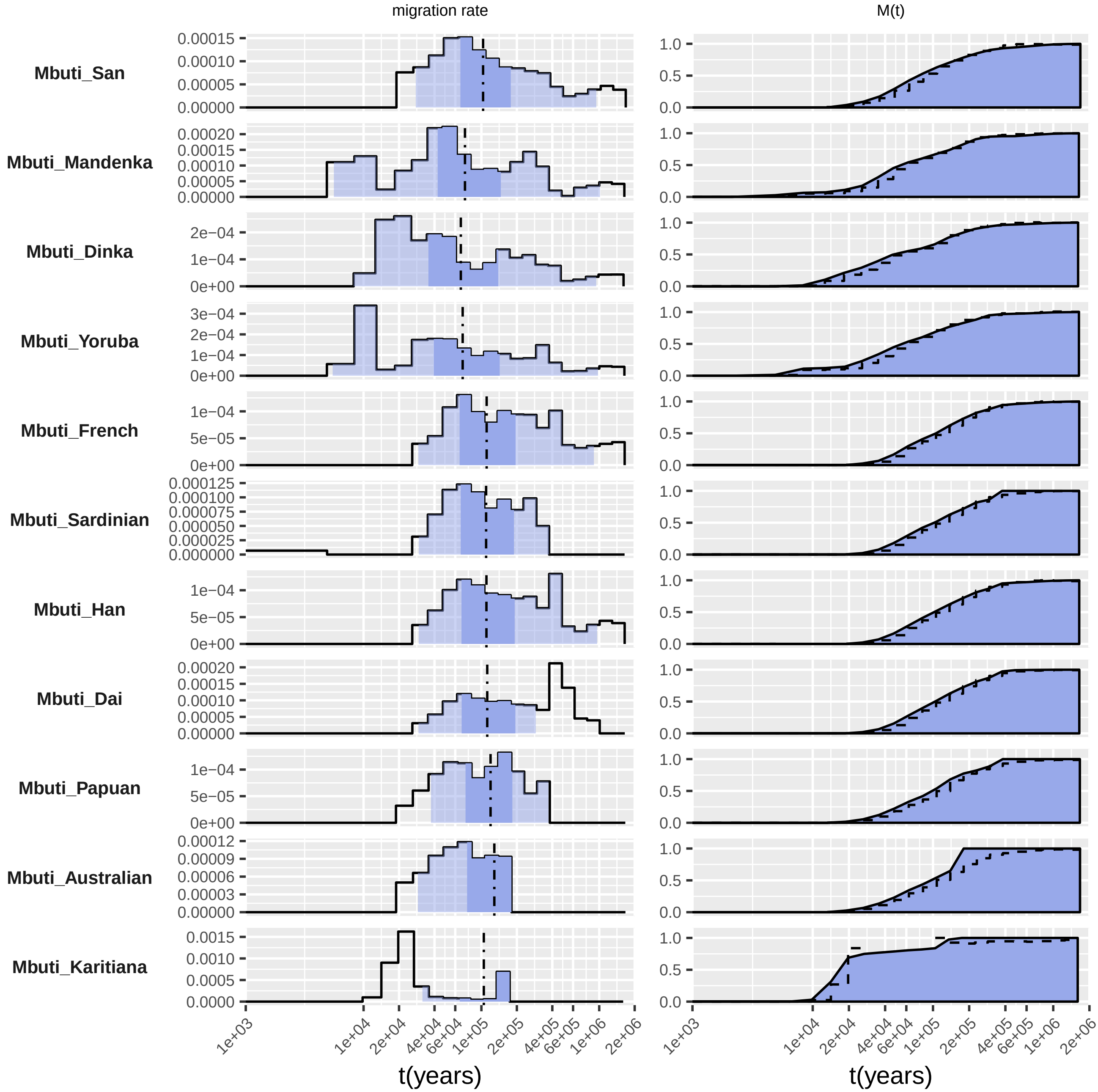

migration rate
M(t)
Mbuti_San
Mbuti_Mandenka
Mbuti_Dinka
Mbuti_Yoruba
Mbuti_French
Mbuti_Sardinian
Mbuti_Han
Mbuti_Dai
Mbuti_Papuan
Mbuti_Australian
Mbuti_Karitiana
1e+03
1e+04
2e+04
4e+04
6e+04
1e+05
2e+05
4e+05
6e+05
1e+06
2e+06
t(years)
t(years)

C

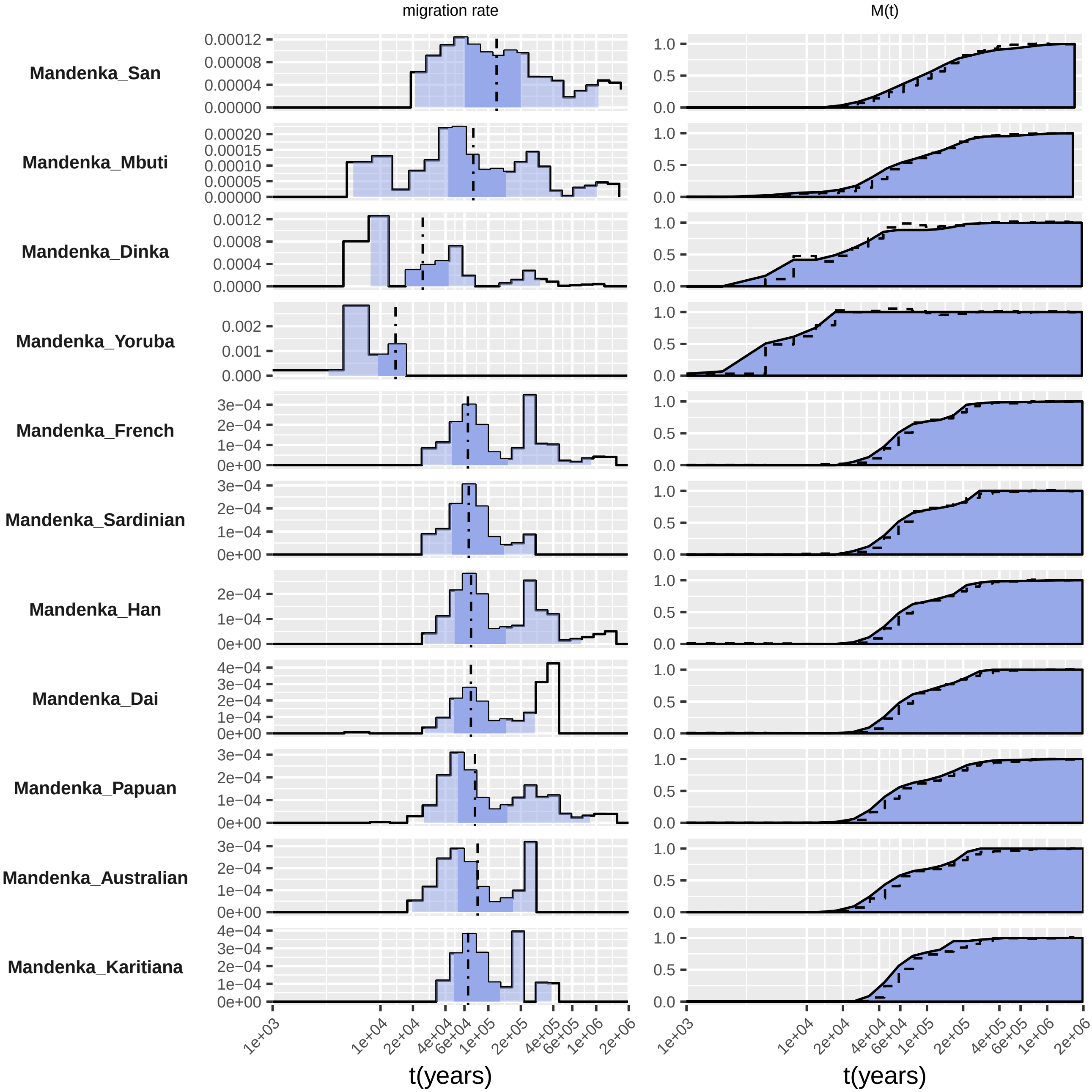

migration rate
M(t)
Mandenka_San
Mandenka_Mbuti
Mandenka_Dinka
Mandenka_Yoruba
Mandenka_French
Mandenka_Sardinian
Mandenka_Han
Mandenka_Dai
Mandenka_Papuan
Mandenka_Australian
Mandenka_Karitiana
1e+03
1e+04
2e+04
4e+04
6e+04
1e+05
2e+05
4e+05
6e+05
1e+06
2e+06
t(years)

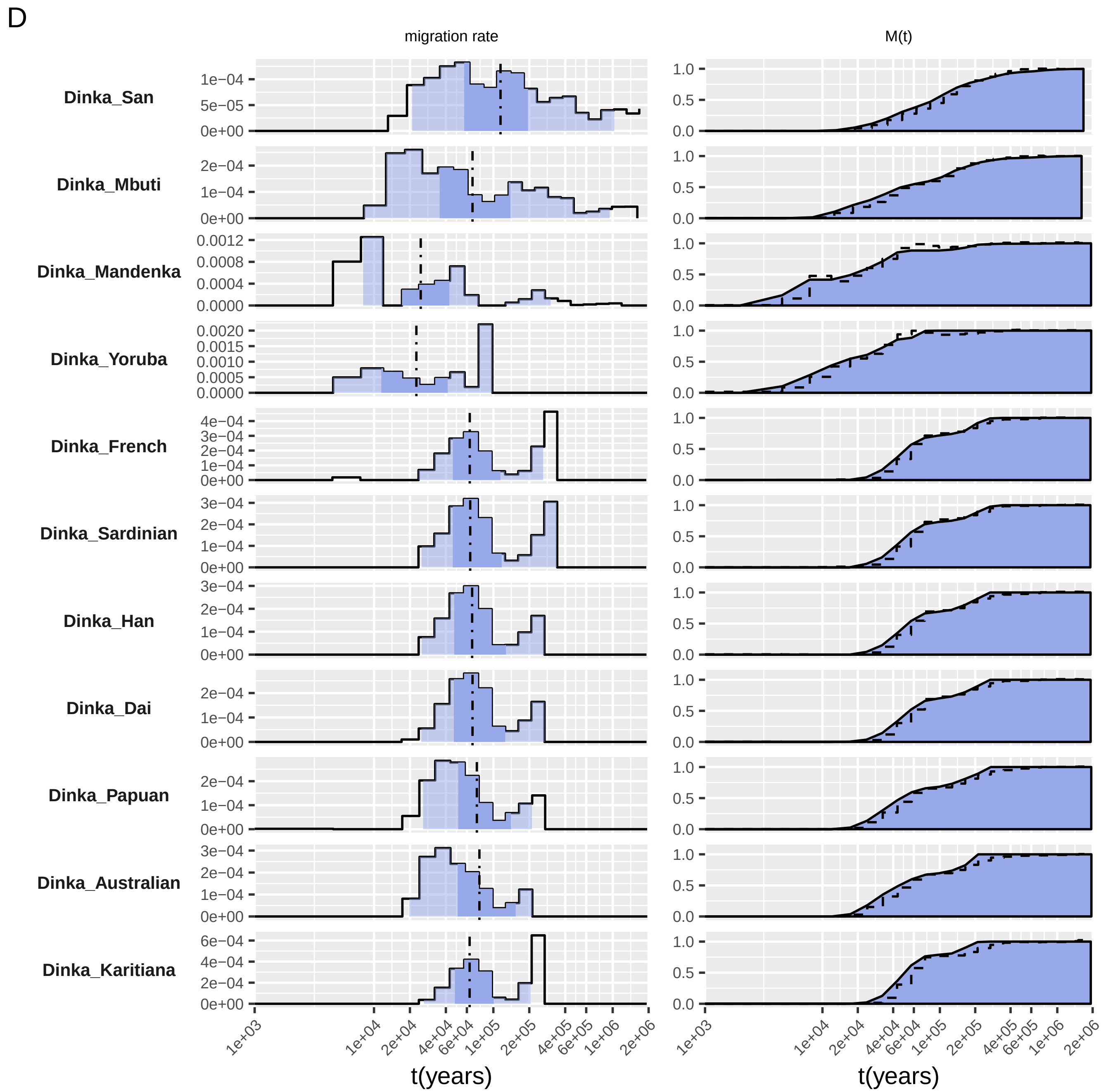

D
migration rate
M(t)
Dinka_San
Dinka_Mbuti
Dinka_Mandenka
Dinka_Yoruba
Dinka_French
Dinka_Sardinian
Dinka_Han
Dinka_Dai
Dinka_Papuan
Dinka_Australian
Dinka_Karitiana
1e+03 1e+04 2e+04 4e+04 6e+04 1e+05 2e+05 4e+05 6e+05 1e+06 2e+06
t(years)
t(years)

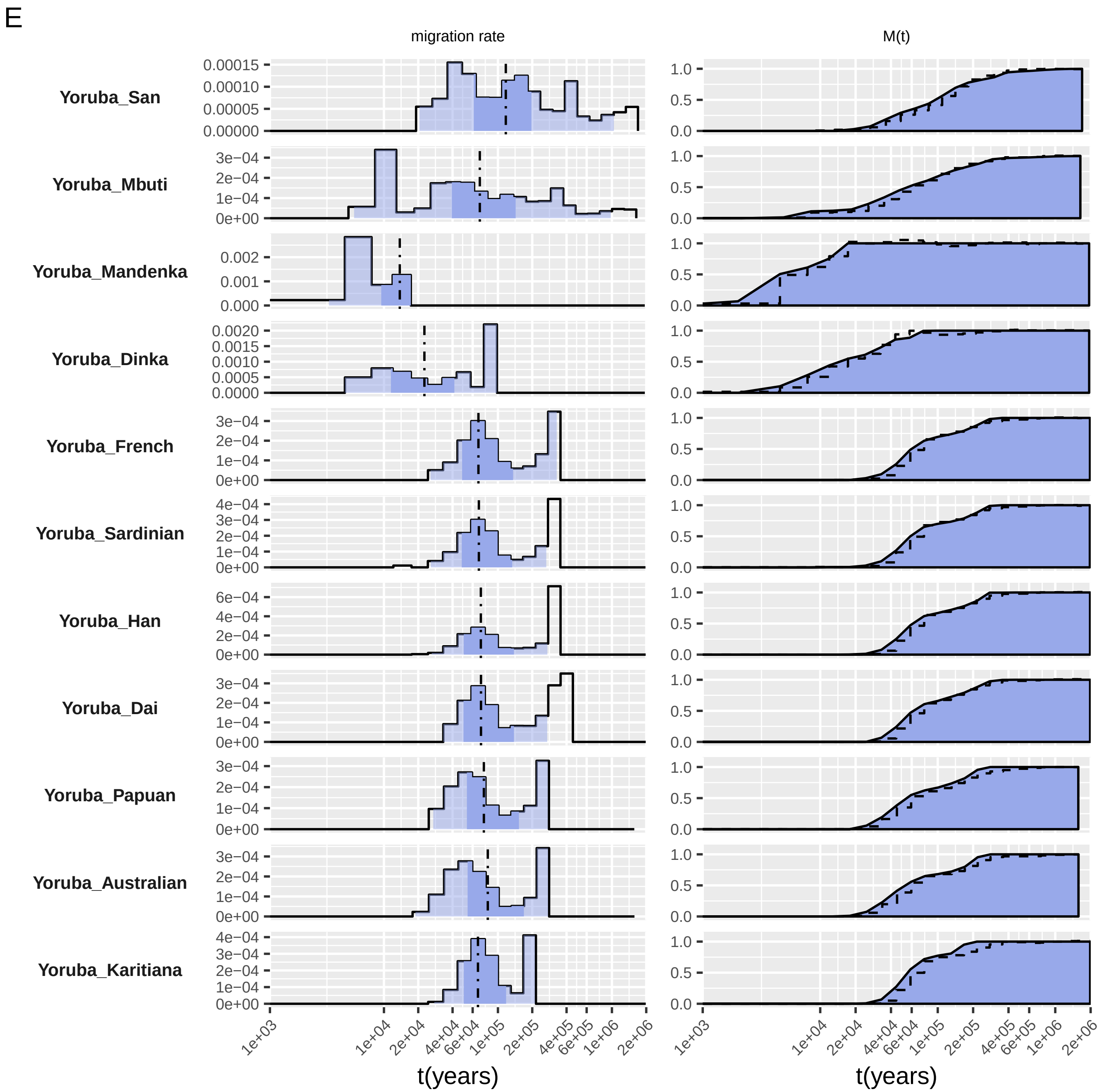
E
migration rate
M(t)
Yoruba_San
Yoruba_Mbuti
Yoruba_Mandenka
Yoruba_Dinka
Yoruba_French
Yoruba_Sardinian
Yoruba_Han
Yoruba_Dai
Yoruba_Papuan
Yoruba_Australian
Yoruba_Karitiana
t(years)
t(years)

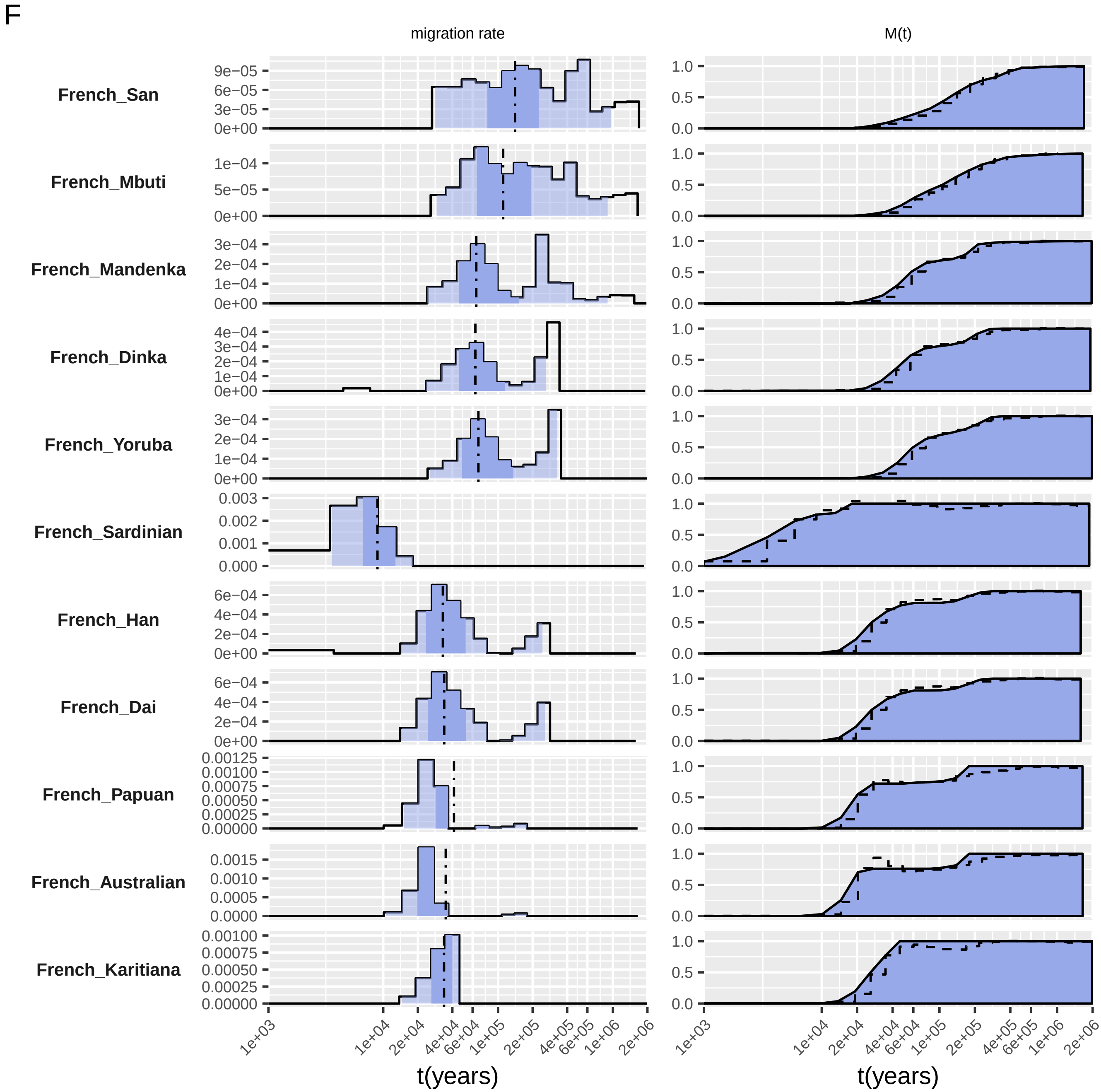
F
migration rate
M(t)
French_San
French_Mbuti
French_Mandenka
French_Dinka
French_Yoruba
French_Sardinian
French_Han
French_Dai
French_Papuan
French_Australian
French_Karitiana
t(years)
t(years)

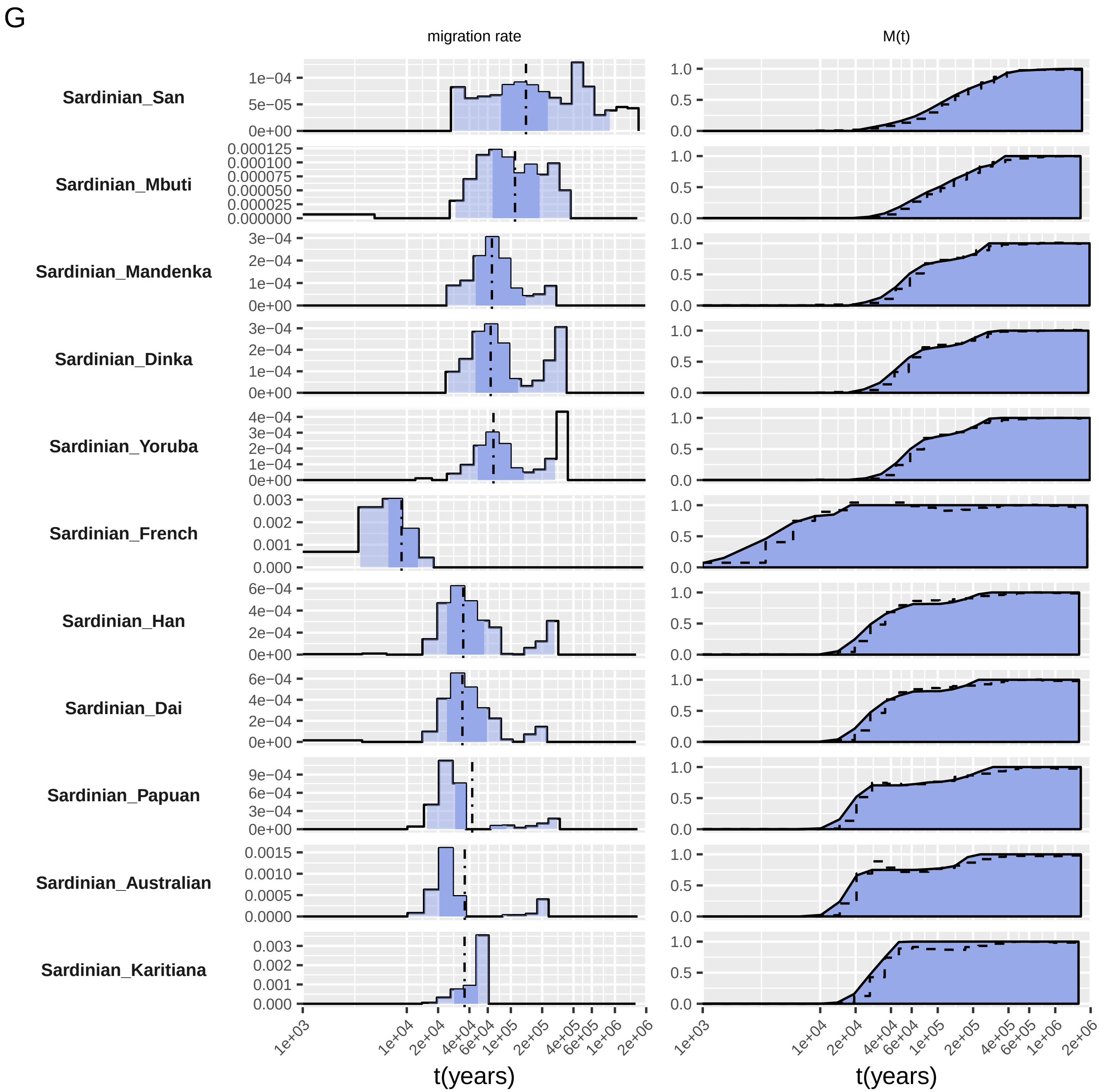

G
migration rate
M(t)
Sardinian_San
Sardinian_Mbuti
Sardinian_Mandenka
Sardinian_Dinka
Sardinian_Yoruba
Sardinian_French
Sardinian_Han
Sardinian_Dai
Sardinian_Papuan
Sardinian_Australian
Sardinian_Karitiana
1e+03
1e+04
2e+04
4e+04
6e+04
1e+05
2e+05
4e+05
6e+05
1e+06
2e+06
t(years)

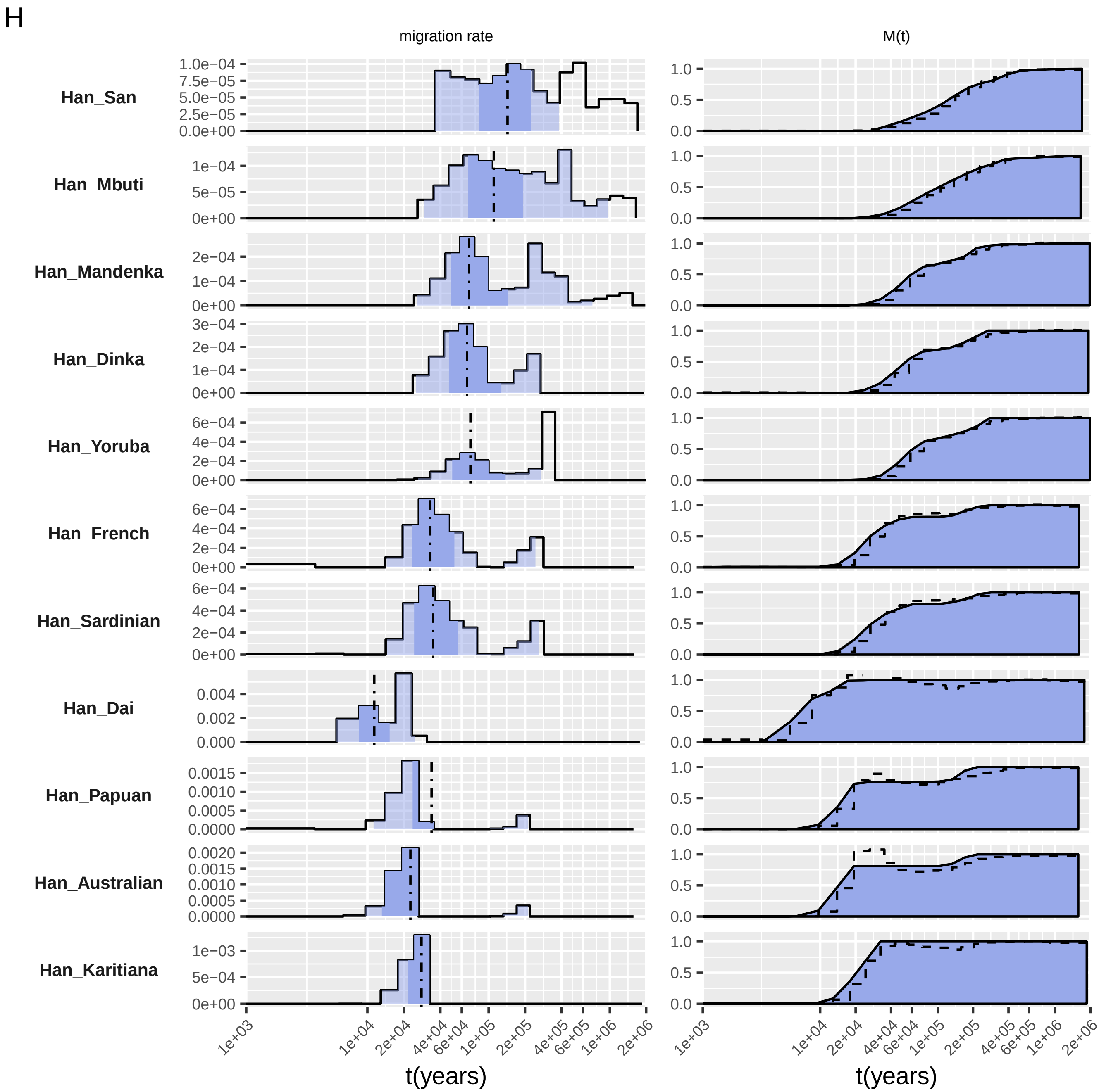
H
migration rate
M(t)
Han_San
Han_Mbuti
Han_Mandenka
Han_Dinka
Han_Yoruba
Han_French
Han_Sardinian
Han_Dai
Han_Papuan
Han_Australian
Han_Karitiana
1e+03
1e+04
2e+04
4e+04
6e+04
1e+05
2e+05
4e+05
6e+05
1e+06
2e+06
t(years)

l

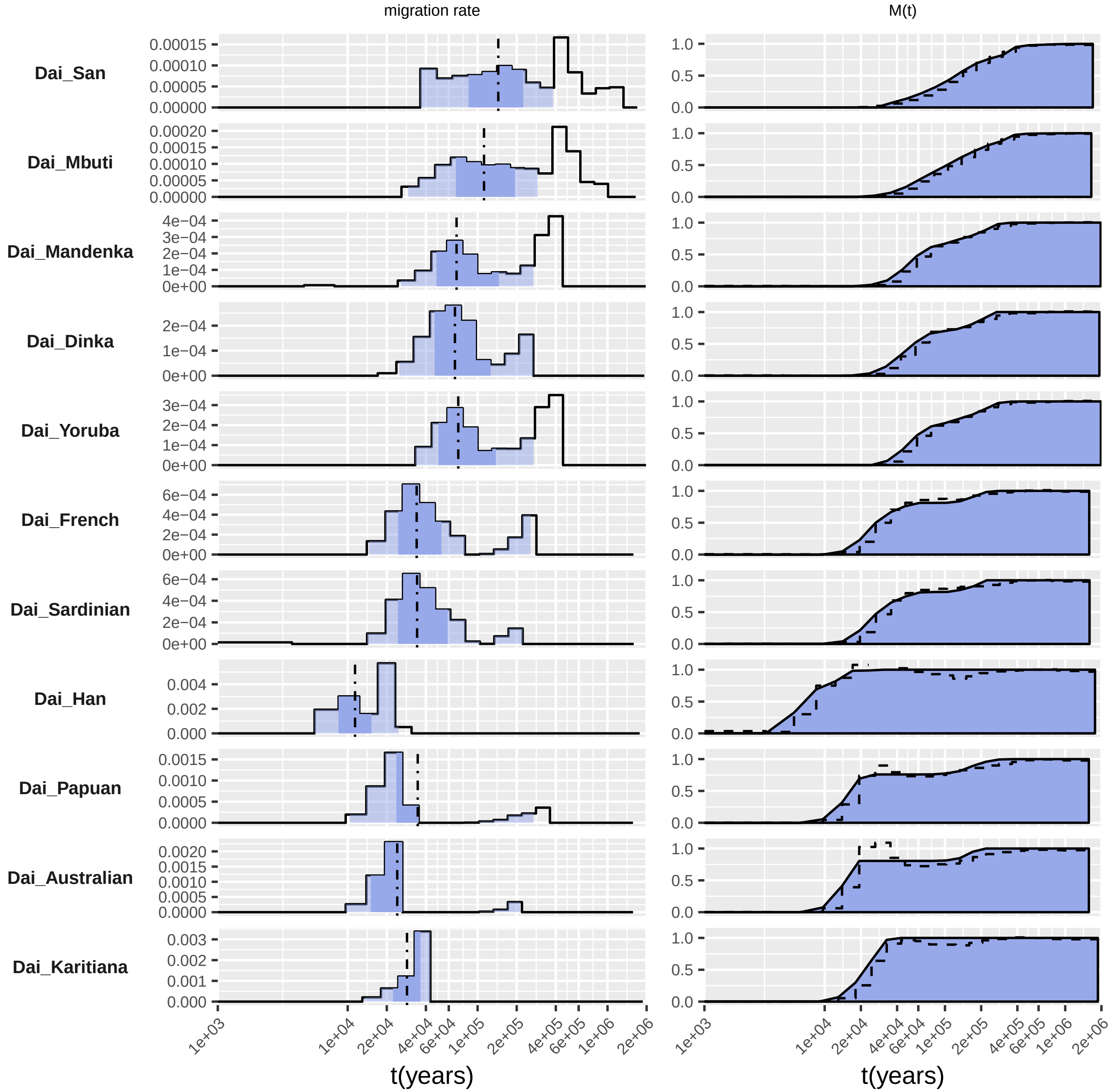

migration rate
M(t)
Dai_San
Dai_Mbuti
Dai_Mandenka
Dai_Dinka
Dai_Yoruba
Dai_French
Dai_Sardinian
Dai_Han
Dai_Papuan
Dai_Australian
Dai_Karitiana
1e+03
1e+04
2e+04
4e+04
6e+04
1e+05
2e+05
4e+05
6e+05
1e+06
2e+06
t(years)
t(years)

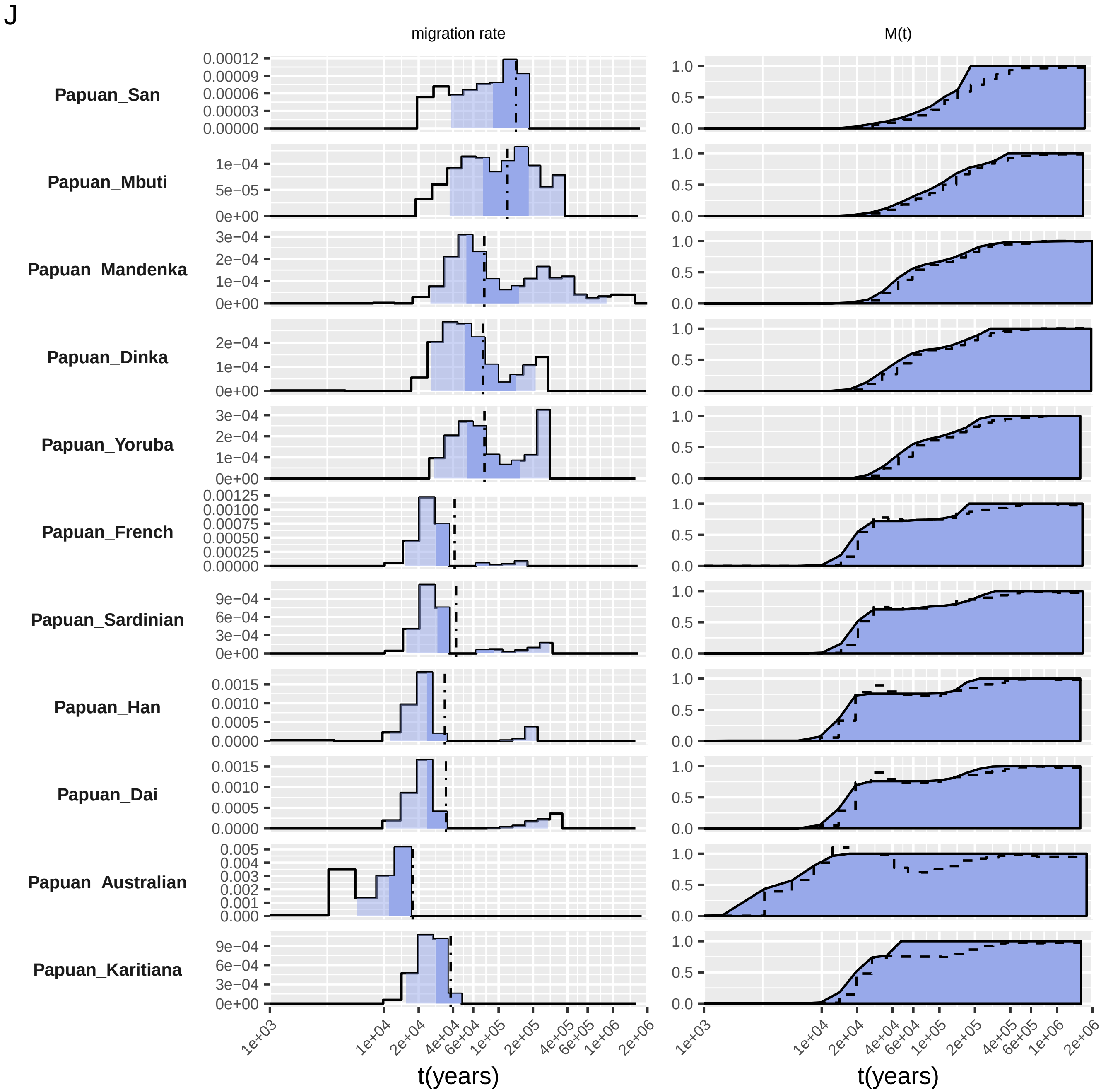

J
migration rate
M(t)
Papuan_San
Papuan_Mbuti
Papuan_Mandenka
Papuan_Dinka
Papuan_Yoruba
Papuan_French
Papuan_Sardinian
Papuan_Han
Papuan_Dai
Papuan_Australian
Papuan_Karitiana
1e+03
1e+04
2e+04
4e+04
6e+04
1e+05
2e+05
4e+05
6e+05
1e+06
2e+06
t(years)

K

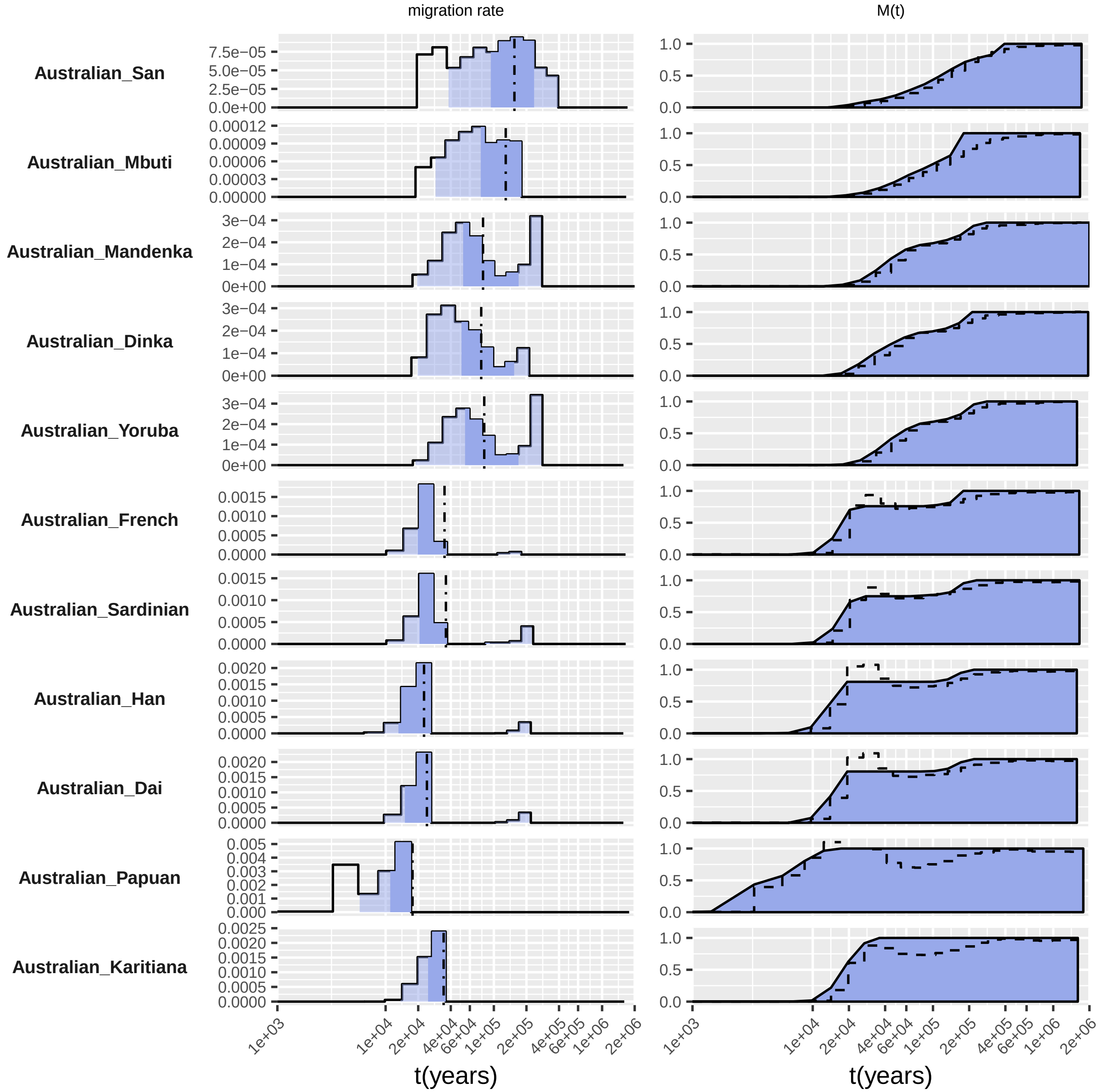

migration rate
M(t)
Australian_San
Australian_Mbuti
Australian_Mandenka
Australian_Dinka
Australian_Yoruba
Australian_French
Australian_Sardinian
Australian_Han
Australian_Dai
Australian_Papuan
Australian_Karitiana
1e+03
1e+04
2e+04
4e+04
6e+04
1e+05
2e+05
4e+05
6e+05
1e+06
2e+06
t(years)

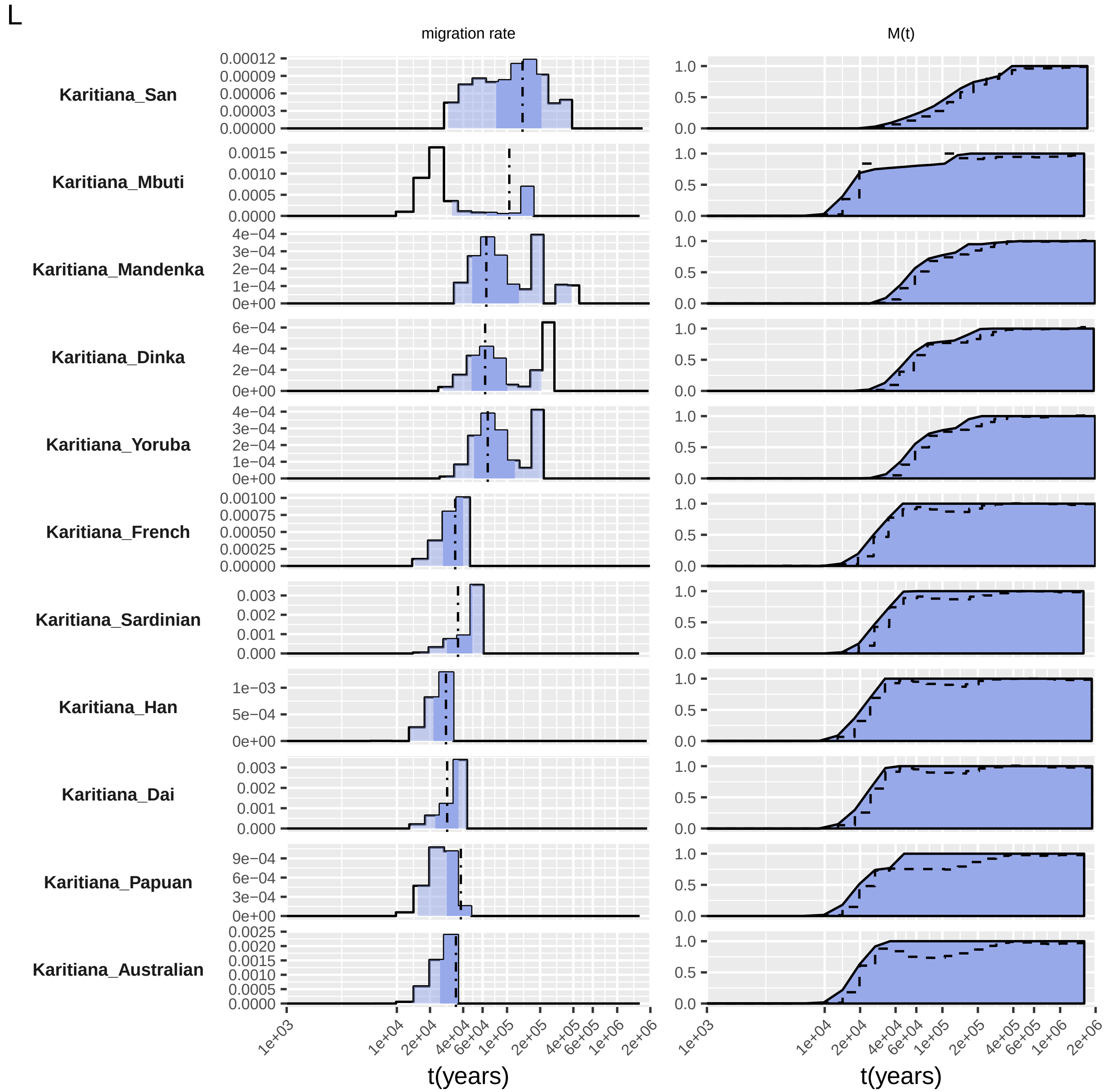
L
migration rate
M(t)
Karitiana_San
Karitiana_Mbuti
Karitiana_Mandenka
Karitiana_Dinka
Karitiana_Yoruba
Karitiana_French
Karitiana_Sardinian
Karitiana_Han
Karitiana_Dai
Karitiana_Papuan
Karitiana_Australian
t(years)
t(years)
