## Supplementary Text for "Tracking human population structure through time from whole genome sequences"

### Supplementary Note for the article "Tracking deep human population structure through time from whole genome sequences"

Ke Wang and Stephan Schiffels

#### 1 MSMC2

MSMC, introduced first in [7] was based on a Hidden Markov Model (HMM) to model the first coalescence event in any two haplotypes in multiple individuals. This approach improved resolution in recent time over PSMC, while sacrificing resolution in ancient times. The newer development MSMC2, first implemented and used in [4], uses a model that is simpler than the HMM of MSMC, and at the same time more powerful. The idea is to run a two-haplotype HMM (called PSMC') on all pairs in a set of multiple haplotypes. The likelihood of the entire data is then multiplied as a composite likelihood. The basic PSMC'-HMM uses only pairs of sequences and hence models only a single coalescence time across a pair of sequences. PSMC' is very similar to PSMC ([3]), but has a more accurate recombination model, based on SMC' [5].

##### 1.1 PSMC

Here we briefly rederive the central equations of the PSMC [3]. In the following, we denote the rate of coalescence by $\lambda(t) = (2N(t))^{-1}$. The transition probability is derived from the SMC model by McVean and Cardin [6]. We consider a *given* recombination event, which takes place at time $u < s$ in either of the two branches $m = \{1, 2\}$. This recombination event causes a "floating" branch which coalesces back onto the other branch at time $t$. The probability for this is given by the probability that *no* coalescence occured between $u$ and $t$ times the probability that it coalesces exactly at time $t$:

$$q(t|s, u, m) = \lambda(t) \exp\left(-\int_u^t \lambda(\nu) d\nu\right) \Theta(t - u) \qquad (1)$$

where the Heavyside-function is defined as

$$\Theta(t-u) = \begin{cases} 1 & \text{if} t > u \\ 0 & \text{else}) \end{cases} \tag{2}$$

and reflects the fact that the transition probability to switch to time $t$ is zero if $u > t$. We show in the Appendix that this conditional probability is properly normalized, i.e. that $\int_0^\infty q(t|s,u,m)\,dt = 1$ for all given $s$, $u$ and $m$.

We need to integrate out the two unknown variables $u$ and $m$, both with uniform probability. The probability that no recombination occured in either of the two branches of length $s$ is $\exp(-2rs)$. Together this yields:

$$q(t|s) = e^{-2rs}\delta(t-s) + \left(1 - e^{-2rs}\right)\frac{1}{s}\frac{1}{2}\int_0^s \sum_{k=1}^{2} q(t|s,u,m)du. \tag{3}$$

or

$$q(t|s) = e^{-2rs}\delta(t-s) + \left(1 - e^{-2rs}\right)\frac{1}{s}\int_0^{\min(s,t)} \lambda(t)\exp\left(-\int_u^t \lambda(\nu)\,d\nu\right)du. \tag{4}$$

#### 1.2 Including Self-coalescence: PSMC'

Marjoram and Wall [5] realized that there was one particular feature missing from the original SMC formulation. An important rationale behind equation 1 is that the recombining "floating" branch will definitely coalesce with the *other* of the two branches, therefore definitely changing the tMRCA to the new value $s$. However, it is of course possible, that the floating branch will simply coalesce back onto its own branch, therefore resulting in a recombination event that does *not* change the tMRCA.

In order to extend the model to include this self-coalescence, we again consider the probability that the time switches from $s$ to $t$, given some recombination time $u$. We can distinguish two cases: for $t > s$, the transition probability is given by the probability that *no* coalescence occured with either of the two branches$<t$ *and* no coalescence to the single branch between $t$ and $s$. For $s < t$, the transition probability is given by the probability of coalescing to the *other* branch, rather than to the self-branch. Finally, we have a third class of recombination events which result in $t = s$, namely if the floating branch coalesces back onto its own branch before $s$.

The conditional probability then reads

$$q(t|s,u,m) = \delta(t-s)\frac{1}{2}\left(1 - \exp\left(-2\int_u^t \lambda(\nu)\,d\nu\right)\right) + \begin{cases} \lambda(t)\exp\left(-\int_u^t 2\lambda(\nu)d\nu\right)\Theta(t-u) & \text{for } t \le s \\ \lambda(t)\exp\left(-\int_u^s 2\lambda(\nu)d\nu - \int_s^t \lambda(\nu)\,d\nu\right) & \text{for } t > s. \end{cases} \tag{5}$$

Again, we show in the Appendix, that this conditional probability is normalized. The full transition probability then reads

$$q(t|s) = \delta(t{-}s)\left(e^{-2rs} + \left(1-e^{-2rs}\right)\frac{1}{2s}\int_0^t \left(1-\exp\left(-2\int_u^t \lambda(\nu)\,d\nu\right)\right)\,du\right) +$$
$$\left(1-e^{-2rs}\right)\frac{1}{s}\begin{cases}\int_0^t \lambda(t)\exp\left(-\int_u^t 2\lambda(\nu)d\nu\right)du & \text{for } t \le s\\ \int_0^s \lambda(t)\exp\left(-\int_u^s 2\lambda(\nu)d\nu - \int_s^t \lambda(\nu)\,d\nu\right)du & \text{for } t > s.\end{cases} \tag{6}$$

The equilibrium probability is

$$q_0(t) = \lambda(t)L(0;t) \tag{7}$$

with the integral

$$L\left(t_1;t_2\right) = \exp\left(-\int_{t_1}^{t_2}\lambda(\nu)\,d\nu\right). \tag{8}$$

For later purposes, we introduce some more functions. We rewrite the transition matrix

$$q(t|s) = \delta(t-s)q_1(t) + q_2(t|s) \tag{9}$$

with

$$q_1(t) = e^{-2rt} + \left(1-e^{-2rt}\right)\frac{1}{2s}\int_0^t \left(1-L(u;t)^2\right)\,du \tag{10}$$

$$q_2(t|s)|_{t<s} = \left(1-e^{-2rs}\right)\frac{1}{s}\lambda(t)\int_0^t L(u;t)^2du, \tag{11}$$

$$q_2(t|s)|_{t>s} = \left(1-e^{-2rs}\right)\frac{1}{s}\lambda(t)L(s;t)\int_0^s L(u;s)^2du. \tag{12}$$

###### 1.2.1 Discrete time intervals

We divide time into a set of $n_T$ intervals that span the entire space from 0 to $\infty$. In practice, as interval boundaries we use the same boundaries as chosen by PSMC [3], defined as:

$$T_i = \alpha\exp\left(\frac{i}{N_T}\log\left(1+\frac{T_{\max}}{\alpha}\right)-1\right) \tag{13}$$

Here, $\alpha$ and $T_{\max}$ are constants that in the case of PSMC were chosen to be $\alpha = 0.1$ and $t_{\max} = 15$. Note that by construction we have $T_0 = 0$ and $T_{N_T} = \infty$.

This patterning sets of with time patterns approximately linearly distributed through time, and then crosses over to a patterning that is uniformly distributed in log-space. This ensures higher resolution in recent than in ancient times.

For MSMC2, we would like to increase resolution in recent times depending on the number of individuals, i.e. haplotypes we use. For example, with four haplotypes, in recent times we have approximately 6 times more recent coalescent events to analyse compared to just two haplotypes . This should therefore allow us to increase resolution in recent times by 6 fold. We generally set the parameter $\alpha$ in equation 13 to be

$$\alpha = \frac{0.1}{n_{\text{pairs}}} \tag{14}$$

where $n_{\text{pairs}}$ is the number of total haplotype pairs analysed. With phased data, and $n_{\text{hap}}$ haplotypes from the same population, we have

$$n_{\text{pairs}} = \frac{n_{\text{hap}}(n_{\text{hap}} - 1)}{2} \tag{15}$$

but this can be different if multiple populations or unphased data is anlaysed. For example, if we have four diploid individuals in total, separated evenly into two populations, then we consider all pairs of haplotypes across the two populations, so we have $n_{\text{pairs}} = 16$. If eight diploid individuals from the same population are analysed, and no phasing is available, then we have $n_{\text{pairs}} = 8$. In the MSMC2-implementation, this behaviour can be controlled with the `--pairIndices` flag (see `https://github.com/stschiff/msmc2`). The scaling of $\alpha$, according to equation 14, is then set automatically by the number of specified pairs.

#### 1.3 Piecewise constant Population sizes

We then define piecewise constant population sizes which correspond to piecewise constant coalescence rates:

$$\lambda(t) = \lambda_\alpha \text{ for } T_\alpha \leq t < T_{\alpha+1}. \tag{16}$$

We now can compute the integral $L(t_1; t_2)$. Let the next *lower* time boundary from $t_1$ be $\beta$, and the next *lower* time boundary from $t_2$ be $\alpha$. We also define $\Delta_\alpha = T_{\alpha+1} - T_\alpha$:

$$L(t_1; t_2)\,|_{\alpha\neq\beta} = \exp\left(-\left(T_{\beta+1} - t_1\right)\lambda_\beta - \sum_{\kappa=\beta+1}^{\alpha-1} \lambda_\kappa \Delta_\kappa - \left(t_2 - T_\alpha\right)\lambda_\alpha\right). \tag{17}$$

$$L(t_1; t_2)\,|_{\alpha=\beta} = \exp\left(-\left(t_2 - t_1\right)\lambda_\alpha\right). \tag{18}$$

In the following, we denote the next lower index of a given time in the function parameters, with $q_0(t;\alpha)$ meaning that $T_\alpha < t < T_{\alpha+1}$:

$$q_0(t;\alpha) = \lambda_\alpha L(0;t) \tag{19}$$

$$q_1(t;\alpha) = e^{-2rt} + \left(1 - e^{-2rt}\right)\frac{1}{2t}\int_0^t \left(1 - L(u;t)^2\right) du \tag{20}$$

For the off-diagonal integrals we first get for $t < s$:

$$q_2(t;\alpha|s)|_{t<s} = \left(1 - e^{-2rs}\right) \frac{1}{s}\lambda(t) \int_0^t L(u;t)^2 du \tag{21}$$

For the case $t > s$, things depend on the interval in which $s$ lies, denoted by $\beta$:

$$\begin{aligned} q_2(t;\alpha|s;\beta)|_{t>s} &= \left(1 - e^{-2rs}\right) \frac{1}{s}\lambda(t)L(s;t) \int_0^s L(u;s)^2 du., \\ &= \left(1 - e^{-2rs}\right) \frac{1}{s}\lambda_\alpha L(s;t) \left( \sum_{\gamma=0}^{\beta-1} \int_{T_\gamma}^{T_{\gamma+1}} L(u;s)^2\, du + \int_{T_\beta}^{s} L(u;s)^2\, du \right) \end{aligned} \tag{22}$$

#### 1.4 Integrating over time intervals

For each time interval we now have to integrate $t$ through $[T_a; T_{a+1}]$. First the equilibrium probability:

$$\begin{aligned} q_0(\alpha) &= \int_{T_\alpha}^{T_{\alpha+1}} \lambda_\alpha L(0;t) dt \\ &= \int_{T_\alpha}^{T_{\alpha+1}} \lambda_\alpha L\left(0;T_\alpha\right) e^{-(t-T_\alpha)\lambda_\alpha} dt \\ &= L\left(0;T_\alpha\right) \left(1 - e^{-\Delta_\alpha \lambda_\alpha}\right) \end{aligned} \tag{23}$$

Next, we compute the expected time in interval $\beta$:

$$\langle t_\beta \rangle = \frac{1}{q_0(\beta)} \int_{T_\beta}^{T_{\beta+1}} t q_0(t;\beta) dt = \frac{1}{L\left(0;T_\beta\right)\left(1 - e^{-\Delta_\beta \lambda_\beta}\right)} \int_{T_\beta}^{T_{\beta+1}} t \lambda_\beta L(0;t) dt \tag{24}$$

This expression for $\langle t_\beta \rangle$ has a numerical instability for $\lambda_\beta \lesssim 10^{-3}$. We set the following asymptotic values:

$$\langle t_\beta \rangle = \begin{cases} \left(T_\beta + T_{\beta+1}\right)/\,2 & \text{for } \lambda_\beta < 10^{-3} \text{ and } T_{\beta+1} < \infty \\ T_\beta + {\lambda_\beta}^{-1} & \text{for } \lambda_\beta < 10^{-3} \text{ and } T_{\beta+1} = \infty \end{cases} \tag{25}$$

We can now write down equations for the off-diagonal elements of the transition matrix, i.e. elements with $\alpha \neq \beta$. First the case $\alpha < \beta$:

$$
\begin{aligned}
q_2(\alpha|\beta)|_{\alpha<\beta} &= \int_{T_\alpha}^{T_{\alpha+1}} q_2\left(t;\alpha|\langle t_\beta\rangle;\beta\right)|_{t<s} dt \\
&= \int_{T_\alpha}^{T_{\alpha+1}} \left(1-e^{-2r\langle t_\beta\rangle}\right)\frac{1}{\langle t_\beta\rangle}\lambda_\alpha \times \\
&\quad \left(\left(\sum_{\gamma=0}^{\alpha-1} L\left(T_{\gamma+1};t\right)^2 \frac{1}{2\lambda_\gamma}\left(1-e^{-2\lambda_\gamma\Delta_\gamma}\right)\right)+\frac{1}{2\lambda_\alpha}\left(1-e^{-2\lambda_\alpha(t-T_\alpha)}\right)\right) dt \\
&= \left(1-e^{-2r\langle t_\beta\rangle}\right)\frac{1}{\langle t_\beta\rangle}\lambda_\alpha\left(\left(1-e^{-2\Delta_\alpha\lambda_\alpha}\right)\sum_{\gamma=0}^{\alpha-1}\left(\frac{1}{2\lambda_\gamma}\left(1-e^{-2\lambda_\gamma\Delta_\gamma}\right)L\left(T_{\gamma+1};T_\alpha\right)^2\right)+\right. \\
&\qquad \left.\frac{1}{2\lambda_\alpha}\left(\Delta_\alpha-\frac{1}{2\lambda_\alpha}\left(1-e^{-2\Delta_\alpha\lambda_\alpha}\right)\right)\right)
\end{aligned} \tag{26}
$$

where we have used

$$
\int_{T_\alpha}^{T_{\alpha+1}} e^{-2(t-T_\alpha)\lambda_\alpha}\,dt = \frac{1}{2\lambda_\alpha}\left(1-e^{-2\Delta_\alpha\lambda_\alpha}\right) \tag{27}
$$

Analogously we have:

$$
\begin{aligned}
q_2(\alpha|\beta)|_{\alpha>\beta} &= \int_{T_\alpha}^{T_{\alpha+1}} q_2\left(t;\alpha\langle t_\beta\rangle;\beta\right)|_{t>s} dt \\
&= \int_{T_\alpha}^{T_{\alpha+1}} \left(1-e^{-2r\langle t_\beta\rangle}\right)\frac{1}{\langle t_\beta\rangle}\lambda_\alpha L\left(\langle t_\beta\rangle;t\right)\times \\
&\quad \left(\sum_{\gamma=0}^{\beta-1}\left(L\left(T_{\gamma+1};\langle t_\beta\rangle\right)^2\frac{1}{2\lambda_\gamma}\left(1-e^{-2\lambda_\gamma\Delta_\gamma}\right)\right)+\frac{1}{2\lambda_\beta}\left(1-e^{-2\lambda_\beta(\langle t_\beta\rangle-T_\beta)}\right)\right) dt \\
&= \int_{T_\alpha}^{T_{\alpha+1}} L\left(\langle t_\beta\rangle;t\right)dt\left(1-e^{-2r\langle t_\beta\rangle}\right)\frac{1}{\langle t_\beta\rangle}\lambda_\alpha\times \\
&\quad \left(\sum_{\gamma=0}^{\beta-1}\left(L\left(T_{\gamma+1};\langle t_\beta\rangle\right)^2\frac{1}{2\lambda_\gamma}\left(1-e^{-2\lambda_\gamma\Delta_\gamma}\right)\right)+\frac{1}{2\lambda_\beta}\left(1-e^{-2\lambda_\beta(\langle t_\beta\rangle-T_\beta)}\right)\right) \\
&= L\left(\langle t_\beta\rangle;T_\alpha\right)\frac{1}{\lambda_\alpha}\left(1-e^{-\Delta_\alpha\lambda_\alpha}\right)\left(1-e^{-2r\langle t_\beta\rangle}\right)\frac{1}{\langle t_\beta\rangle}\lambda_\alpha\times \\
&\quad \left(\sum_{\gamma=0}^{\beta-1}\left(L\left(T_{\gamma+1};\langle t_\beta\rangle\right)^2\frac{1}{2\lambda_\gamma}\left(1-e^{-2\lambda_\gamma\Delta_\gamma}\right)\right)+\frac{1}{2\lambda_\beta}\left(1-e^{-2\lambda_\beta(\langle t_\beta\rangle-T_\beta)}\right)\right)
\end{aligned} \tag{28}
$$

where we have used

$$\int_{T_\alpha}^{T_{\alpha+1}} L(s;t) = L\left(s;T_\alpha\right) \int_{T_\alpha}^{T_{\alpha+1}} e^{-(t-T_\alpha)\lambda_\alpha}\, dt = L\left(s;T_\alpha\right) \frac{1}{\lambda_\alpha}\left(1-e^{-\Delta_\alpha\lambda_\alpha}\right) \tag{29}$$

The complete discrete transition matrix now reads:

$$q(\alpha|\beta) = \delta_{\alpha,\beta} q_1(\beta) + q_2(\alpha|\beta) \tag{30}$$

with

$$q_1(\beta) = 1 - \sum_{\alpha\neq\beta} q_2(\alpha|\beta) \tag{31}$$

due to the column normalization of the transition matrix.

#### 1.5 Emission Probability

An observation at location $i$ in the genome for a pair of haplotypes (as in a single diploid genome), $O_i$, can be either of $O_i = \{0, 1, 2\}$, where 0 denotes missing data in either of the two haplotypes, 1 denotes a site where both haplotypes have the same allele (i.e. a homozygous genotype in case of a single diploid genome), 1 denotes a mismatch between the alleles of the two haplotypes (i.e. a heterozygote genotype in case of a single diploid genome),

The emission probabilities for exact coalescence times are:

$$e(0|t) = 1 \tag{32}$$
$$e(1|t) = e^{-2\mu t} \tag{33}$$
$$e(2|t) = 1 - e(1|t) \tag{34}$$

For discrete time intervals, we we need to integrate over the conditional probability distribution in each time interval:

$$\begin{aligned}
e(0|\alpha) &= 1 \\
e(1|\alpha) &= \frac{\int_{T_\alpha}^{T_{\alpha+1}} q_0(t) e^{-2\mu t} dt}{\int_{T_\alpha}^{T_{\alpha+1}} q_0(t)\, dt} = \frac{\int_{T_\alpha}^{T_{\alpha+1}} \lambda_\alpha L(0;t) e^{-2\mu t} dt}{L\left(0;T_\alpha\right)\left(1-e^{-\Delta_\alpha\lambda_\alpha}\right)} \\
&= \frac{\lambda_\alpha}{\left(1-e^{-\Delta_\alpha\lambda_\alpha}\right)} \int_{T_\alpha}^{T_{\alpha+1}} L\left(T_\alpha;t\right) e^{-2\mu t} dt \\
&= \frac{\lambda_\alpha}{\left(1-e^{-\Delta_\alpha\lambda_\alpha}\right)} \int_{T_\alpha}^{T_{\alpha+1}} e^{-(t-T_\alpha)\lambda_\alpha} e^{-2\mu t} dt \\
&= \frac{\lambda_\alpha e^{T_\alpha\lambda_\alpha}}{\left(1-e^{-\Delta_\alpha\lambda_\alpha}\right)} \int_{T_\alpha}^{T_{\alpha+1}} e^{-(2\mu+\lambda_\alpha)t} dt \\
&= \frac{\lambda_\alpha e^{T_\alpha\lambda_\alpha}}{\left(1-e^{-\Delta_\alpha\lambda_\alpha}\right)} \frac{e^{-2\mu T_\alpha}}{2\mu+\lambda_\alpha} \left(1-e^{-(2\mu+\lambda_\alpha)\Delta_\alpha}\right)
\end{aligned} \tag{35}$$

and of course we have as before:

$$e(2|\alpha) = 1 - e(1|\alpha) \tag{36}$$

There are special forms of these expressions for two cases. First, if $T_{\alpha+1} = \infty$, then we have $\Delta_\alpha = \infty$, and so the expression becomes

$$e(1|\alpha)|_{T_{\alpha+1}=\infty} = \lambda_\alpha \frac{e^{-2\mu T_\alpha}}{2\mu + \lambda_\alpha} \tag{37}$$

Second, there is again a numerical instability for $\lambda_\alpha \lesssim 10^{-3}$, in which case the expression becomes

$$e(1|\alpha)|_{\lambda_\alpha \lesssim 10^{-3}} = \frac{1}{2\Delta_\alpha \mu} e^{-2\mu T_\alpha} \left(1 - e^{-(2\mu+\lambda_\alpha)\Delta_\alpha}\right) \tag{38}$$

#### 1.6 MSMC2 Hidden Markov Model

We can now define a Hidden Markov Model (see [1] for background reading), based on PSMC' using the above defined transition and emission probabilities. For a given sequence of length $L$, we define the observations as $O_1 \dots O_L$. We define a forward variable $f_1(\alpha) \dots f_L(\alpha)$ by the recursion relation:

$$f_1(\alpha) = q_0(\alpha) e(O_1|\alpha) \tag{39}$$

$$f_n(\alpha) = e(O_n|\alpha) \sum_\beta q(\alpha|\beta) f_{n-1}(\beta) \quad \text{for } n = 2 \dots L \tag{40}$$

Analogously, a "backwards"-vector $b_1(\alpha) \dots b_L(\alpha)$ is defined as:

$$b_L(\alpha) = 1 \tag{41}$$

$$b_n(\beta) = \sum_\alpha e(O_{n+1}|\alpha) q(\alpha|\beta) b_{n+1}(\alpha) \quad \text{for } n = (L-1) \dots 1 \tag{42}$$

In practice, we can speed these algorithms up substantially by precomputing powers of emission-transition matrices in order to quickly skip over long regions with missing or homozygous data. This is described in [7].

We now recursively run these two variables over all chromosomes and all pairs of haplotypes. This makes it different from MSMC, which consisted of one HMM across all haplotypes simultaneously. Here we run separately over all combinations of pairs. So for example, with two diploid phased human genomes from a single population, we would run the forward-backward algorithm independently (and possibly in parallel) over 132 chromosomal pairs of haplotypes: 6 pairs of haplotypes ((1,2), (1,3), (1,4), (2,3), (2,4), (3,4)) on 22 chromosomes each.

In order to estimate parameters of our HMM (i.e. the piecewise constant coalescence rates $\lambda_\alpha$ and the recombination rate $r$), we use the Baum-Welch algorithm, similarly to MSMC.

We first define an objective function

$$F(\theta,\bar{\theta}) = \sum_{\alpha,\beta} \log(q(\alpha|\beta;\bar{\theta}))\Xi(\alpha|\beta,O_n,\theta) + \sum_{O',\alpha} \log(e(O'|\alpha;\bar{\theta}))\Gamma(O',\alpha;O_n,\theta) \tag{43}$$

with

$$\Xi(\alpha|\beta,O_n,\theta) = \sum_n f_n(\beta)q(\alpha|\beta)e(O_{n+1}|\alpha)b_{n+1}(\alpha), \tag{44}$$

and

$$\Gamma(O'|\alpha,\theta) = \sum_n f_n(\alpha)b_n(\alpha)e(O_n|\alpha)I(O_n = O') \tag{45}$$

where $O_n$ denotes the entire collection of observed data across all chromosomes and analysed haplotype pairs, $\theta$ denotes the set of parameters used in this iteration of the algorithm, $\bar{\theta}$ denotes free parameters to be varied in the maximization step of the algorithm (see below). The first term in equation 43 sums up the evidence from the observed transitions along the data, and the second sums up the evidence from the observed emissions. Both evidence matrices depend on the data and on the current set of parameters $\theta$. Matrix $\Xi$ is a square-matrix with as many rows and columns as there are hidden states. Matrix $\Gamma$ has as many rows as there are different symbols in the alphabet (here 3), and as many columns as there are hidden states.

The sum runs in principle over all sites across all analysed chromosomes and haplotype pairs. In practice, we sparsen this sum by selecting an equally spaced set of sites. By default, the distance between each counted site is 1000, but this can be controlled via the parameter `--hmmStrideWidth`.

The maximization step of the Baum-Welch algorithm then re-estimates the parameters by maximizing the objective function:

$$\hat{\theta} = \arg\max_{\bar{\theta}} F(\theta,\bar{\theta}) \tag{46}$$

The Baum-Welch algorithm consists of iterations of i) the forward-backward algorithm to compute the objective function, and ii) a maximization step to estimate new parameters. In the next iteration, the forward-backward algorithm is then run with the new parameters, and so forth.

After about 20 iterations, we find that the likelihood plateaus for most MSMC runs.

#### 1.7 Combining within- and cross-coalescence rates estimates

While MSMC can estimate three coalescence rate functions simultaneously when run over genomes from two populations, MSMC2 runs over pairs of populations separately. Each run then uses a slightly different time scaling (due to different heterozygosity, i.e. allele mismatch, estimates within and across populations). For MSMC-IM, we however need three estimates of coalescence rates defined along the same time intervals.

We supply a simple python script, called `combineCrossCoal.py`, which reads in three result files from MSMC2, each from one pair of populations, and uses interpolation of the resulting piecewise constant coalescence rate estimates to merge these datasets. Details about this can be found in the accompanying README of the `msmc-tools` repository on `github.com/stschiff/msmc-tools`.

#### 1.8 Appendix: Normalizations

In the following derivations, we define $L(t)$ to be an antiderivative of $\lambda(t)$, i.e. $L'(t) = \lambda(t)$. We will also make use of the substitution rule

$$\int_a^b g'(x)f(g(x))dx = \int_{g(a)}^{g(b)} f(z)\,dz. \tag{47}$$

##### 1.8.1 PSMC conditional transition probability

We have

$$q(t|s,u,m) = \lambda(t)\exp\left(-\int_u^t \lambda(v)\,dv\right)\Theta(t-u). \tag{48}$$

We need to show that the PSMC conditional probability is normalized:

$$\begin{aligned}
\int_0^\infty q(t|s,u,m)\,dt &= \int_0^\infty \lambda(t)\exp\left(-\int_u^t \lambda(\nu)\,d\nu\right)\Theta(t-u)dt \\
&= \int_u^\infty \lambda(t)\exp\left(-\int_u^t \lambda(\nu)\,d\nu\right) \\
&= \int_u^\infty \lambda(t)\exp\left(-\int_u^t \lambda(\nu)\,dv\right) \\
&= \int_u^\infty L'(t)e^{-(L(t)-L(u)} = e^{L(u)}\int_{L(u)}^{L(\infty)} e^{-z}\,dz = e^{L(u)}\left(e^{-L(u)} - e^{-L(\infty)}\right) \\
&= 1-\exp\left(-\int_u^\infty \lambda(\nu)\,d\nu\right) \\
&= 1 \quad \square
\end{aligned} \tag{49}$$

##### 1.8.2 PSMC' conditional transition probability

We have

$$q(t|s,u,m) = \delta(t-s)\frac{1}{2}\left(1-\exp\left(-2\int_u^t \lambda(\nu)\,d\nu\right)\right) + \begin{cases} \lambda(t)\exp\left(-\int_u^t 2\lambda(\nu)dv\right)\Theta(t-u) & \text{for } t \leq s \\ \lambda(t)\exp\left(-\int_u^s 2\lambda(\nu)d\nu - \int_s^t \lambda(\nu)\,d\nu\right) & \text{for } t > s. \end{cases} \tag{50}$$

We again need to compute the integral $\int_0^\infty q(t|s,u,m)\,dt$. We divide the integral into three parts:

$$\begin{aligned}
\int_0^\infty q(t|s,u,m)\,dt &= \int_0^\infty \delta(t-s)\frac{1}{2}\left(1-\exp\left(-2\int_u^t \lambda(v)\,dv\right)\right)dt + \\
&\int_u^s \lambda(t)\exp\left(-\int_u^t 2\lambda(v)dv\right)dt + \\
&\int_s^\infty \lambda(t)\exp\left(-\int_u^s 2\lambda(v)dv - \int_s^t \lambda(v)\,dv\right)dt \\
&= \frac{1}{2}\left(1-e^{-2(L(s)-L(u))}\right) + \int_u^s L'(t)e^{-2(L(t)-L(u))}dt + \\
&e^{-2(L(s)-L(u))}\int_s^\infty L'(t)e^{-(L(t)-L(s))}dt \\
&= \frac{1}{2}\left(1-e^{-2(L(s)-L(u))}\right) + e^{2L(u)}\int_{L(u)}^{L(s)} e^{-2z}\,dz + \\
&e^{-2(L(s)-L(u))}e^{L(s)}\int_{L(s)}^{L(\infty)} e^{-z}\,dz \\
&= \frac{1}{2}\left(1-e^{-2(L(s)-L(u))}\right) + e^{2L(u)}\frac{1}{2}\left(e^{-2L(u)}-e^{-2L(s)}\right) + \\
&e^{-2(L(s)-L(u))}e^{L(s)}\left(e^{-L(s)}-e^{-L(\infty)}\right) \\
&= \frac{1}{2}\left(1-e^{-2(L(s)-L(u))}\right) + \frac{1}{2}\left(1-e^{2(L(u)-L(s))}\right) + e^{-2(L(s)-L(u))}\left(1-e^{L(s)-L(\infty)}\right) \\
&= 1-e^{-2(L(s)-L(u))} + e^{-2(L(s)-L(u))} \\
&= 1 \quad \square
\end{aligned} \tag{51}$$

### 2 MSMC-IM model

#### 2.1 Continuous IM model

Our model is based on Hobolth et al. 2011 [2], which demonstrates that the time to the most recent common ancestor (tMRCA) of two lineages sampled from a pair of populations can be exactly computed from a matrix exponential. Hobolth et al. 2011 [2] formulate the IM model as a continuous time Markov chain.

Here we build on that work and define a two-island model by time-dependent population sizes $N_1(t)$ and $N_2(t)$ and a time-dependent continuous symmetric migration rate $m(t)$ between the two populations, discarding the clean split concept in Hobolth et al. but describe the population separation as a continuous process.

The state space of our Markov chain matches the state space from the model in Hobolth et al. for times more recent than the split time. There are five possible states of uncoalesced and coalesced lineages: $S_{11}$ denotes two uncoalasced lineages residing in population 1; $S_{12}$ denotes the state where one lineage resides in population 1 and the other in population 2; $S_{22}$ denotes both lineages residing in population 2; $S_1$ describes the state where the two lineages have coalesced, and the single remaining lineage resides in population 1; $S_2$ similarly, where the single remaining lineage resides in population 2.

The state of the two lineages composes a series of states in a Markov chain. At time $t = 0$ (the present-day generation), the state of two randomly sampled uncoalesced lineages starts from either of the following three states $S_{11}$, $S_{12}$, $S_{22}$, and at any later time end up in any of five states $S_{11}$, $S_{12}$, $S_{22}$, $S_1$ or $S_2$.

We describe this evolution of the state space via a probability vector $x_n(t)$ denoting the state probability to be in state $n$ at time $t$, with time counting backwards in time. We summarise that vector in bold font as $\mathbf{x}(t)$.

We summarise the transition rate between states by a matrix $\mathbf{Q}(t)$, where rows indicating the state at some time $t$, and columns the state one generation later. Then the matrix $\mathbf{Q}$ can be expressed in terms of a symmetric migration rate and effective population sizes (very similar to [2]):

$$\mathbf{Q} = \begin{array}{c} \\ S_{11} \\ S_{12} \\ S_{22} \\ S_1 \\ S_2 \end{array} \begin{array}{c} \begin{array}{ccccc} S_{11} & S_{12} & S_{22} & S_1 & S_2 \end{array} \\ \begin{pmatrix} \cdot & 2m(t) & 0 & \frac{1}{2N_1(t)} & 0 \\ m(t) & \cdot & m(t) & 0 & 0 \\ 0 & 2m(t) & \cdot & 0 & \frac{1}{2N_2(t)} \\ 0 & 0 & 0 & \cdot & m(t) \\ 0 & 0 & 0 & m(t) & \cdot \end{pmatrix} \end{array}$$

where $N_1(t)$, $N_2(t)$ and $m(t)$ are all time-dependent. Diagonal elements are set such that rows sum up to zero. The state probability vector in the next generation is then the product of $\mathbf{x}(t)$ and $\mathbf{Q}$:

$$\mathbf{x}(t+1) = \mathbf{x}(t) \cdot (\mathbf{1} + \mathbf{Q}), \tag{52}$$

where $\mathbf{1}$ is a diagonal unit matrix. For $n$ generations, we get

$$\mathbf{x}(t+n) = \mathbf{x}(t) \cdot (\mathbf{1} + \mathbf{Q})^n. \tag{53}$$

We now switch to continuous time, and note that for a small time interval $\Delta t$ we can write:

$$\mathbf{x}(t_0 + \Delta t) = x(t_0) \cdot (1 + \Delta t \mathbf{Q}) \tag{54}$$

Longer time segments $t$ can then be divided into $n$ small time intervals, and we assume Q is constant in each interval and independent from matrices in other intervals.

$$\mathbf{x}(t_0 + t) = \mathbf{x}(t_0) \cdot \left(\mathbf{1} + \frac{t}{n}\mathbf{Q}\right)^n \tag{55}$$

In the limit of $n \to \infty$, the equation above becomes a matrix exponential:

$$\mathbf{x}(t_0 + t) = \mathbf{x}(t_0) \cdot e^{\mathbf{Q}t} \tag{56}$$

When $t_0 = 0$, we then have:

$$\mathbf{x}(t) = \mathbf{x}(0) \cdot e^{\mathbf{Q}t} \tag{57}$$

We can use this general state propagation equation to compute the conditional probability of ending up in a specific final state $s_f$ after time t given a specific starting state $s_0$. For example, the probability to end in state $s_f = S_{11}$ when starting in state $s_0 = S_{12}$ would be:

$$G(s_f = S_{11}, t | s_0 = S_{12}) = \left[ \begin{pmatrix} 0 \\ 1 \\ 0 \\ 0 \\ 0 \end{pmatrix} \cdot e^{\mathbf{Q}t} \right]_{S_{11}} \tag{58}$$

where we have followed the convention introduced above that the order of states in vector notation is $S_{11}$, $S_{12}$, $S_{22}$, $S_1$, $S_2$.

We can now use this to write down the probability of a coalescence event of the two lineages at time $t$, starting in on of the starting states $s_0 \in \{S_{11}, S_{12}, S_{22}\}$:

$$\mathbf{P}^{\mathrm{IM}}(t | s_0, N_1, N_2, m) = G(S_{11}, t | s_0) \cdot 1/2N_1 + G(S_{22}, t | s_0) \cdot 1/2N_2 \tag{59}$$

because in order for a coalescence to occur exactly at time $t$, we reuqire that i) no coalescence has occured before (so we exclude final states $S_1$ and $S_2$), ii) both lineages are in the same population (so we exclude $S_{12}$).

#### 2.2 Comparing with MSMC outputs

MSMC (here as a term used independently from a specific implementation like MSMC or MSMC2) estimates time-dependent effective coalescent rates $\lambda_{ij}$ between a pair of lineages $i$ and $j$. From these rates, we can compute the probability density for coalescence events:

$$\mathbf{P}^{\mathrm{MSMC}}(t|s_0 = S_{ij}) = \lambda_{ij}(t) \cdot e^{-\int_0^t \lambda_{ij}(t')dt'} \tag{60}$$

The basic idea behind MSMC-IM is to fit the model from equation 59 to the observed distribution from equation 60 to estimate parameters $N_1(t)$, $N_2(t)$ and $m(t)$.

#### 2.3 Model Fitting

So far we haven't specified the form of the time-dependent parameters $N_1(t)$, $N_2(t)$ and $m(t)$. Since MSMC uses piecewise constant functions for the coalescence rates, we decided to use exactly the same method in MSMC-IM, and impose a piece-wise constant structure on our model parameters with the same time patterning as in MSMC.

We denote the mid-time points for time segment $i$ by $t_i$, with $i = 1 \dots n_T$. We can then define the following $\chi^2$-statistic across all time-segments to measure the fit deviation between the coalescent distributions from MSMC and the IM model:

$$\tilde{\chi}^2 = \sum_{i=1}^{n_T} \sum_{x_0 \in \{S_{11}, S_{12}, S_{22}\}} \frac{(\mathbf{P}^{IM}(t_i|s_0) - \mathbf{P}^{MSMC}(t_i|s_0))^2}{\mathbf{P}^{MSMC}(t_i|s_0)} \tag{61}$$

For brevity we omit the dependency on model parameters $N_1(t)$, $N_2(t)$ and $m(t)$ here. Minimization of this $\chi^2$-statistic is numerically implemented via Powell's method (using the function `minimize(method='Powell')` from the scipy-package in python (www.scipy.org)).

##### 2.3.1 Regularisation

We need to estimate $N_1$, $N_2$, m for each time interval, which for the default MSMC time patterning means 96 parameters in total. To avoid over-fitting, we add a regularisation term to the above $\chi^2$-statistic:

$$\tilde{\chi}^2 = \sum_{i=1}^{n_T} \left[ \sum_{s_0 \in \{S_{11}, S_{12}, S_{22}\}} \frac{(\mathbf{P}^{IM}(t_i|s_0) - \mathbf{P}^{MSMC}(t_i|s_0))^2}{\mathbf{P}^{MSMC}(t_i|s_0)} + \beta \left( \frac{N_1(t) - N_2(t)}{N_1(t) + N_2(t)} \right)^2 \right] \tag{62}$$

The regularization term $\beta$ is tunable, and we in practice set it to 1e-6 by default. This value was chosen to be low enough to not affect population size estimates at time substantially before the split time, but strong enough to "pull together" the two population sizes for times very deep in the past, where all lineages have effectively merged into one population.

##### 2.3.2 Hazard function for estimating coalescence rates from IM model

While the primary variable to use for comparison between model and data is the probability density function of pairwise coalescence times (eqs. 60 and 59), we can also compute the Hazard function from the model, to be directly compared to the pairwise coalescence rates output by MSMC: as following equation:

$$\lambda_{ij}^{IM}(t) = \frac{\mathbf{P}(t|s_0 = S_{ij}, N_1, N_2, m)}{1 - \int_0^t \mathbf{P}(t|s_0 = S_{ij}, N_1, N_2, m)} \tag{63}$$

This expression becomes numerically unstable for very ancient times, for which the denominator becomes too small.

##### 2.3.3 Internal auto-Correction and parameter constraints

In some cases, MSMC coalescence rate estimates in the most ancient few time intervals are noisy, which can affect migration rate estimates in these windows and lead to artifacts. We therefore implemented an automatic check of the rate estimates in the most ancient time intervals before fitting with MSMC-IM, and auto-correct these values. Specifically, we check in all time segments that correspond to the last two free parameters (with the default patterning of 1*2+25*1+1*2+1*3, as in MSMC2, the last five time intervals would be checked). In these intervals, since we do not genuinely expect estimates to fluctuate much at this end of the analysis time window, we require estimates to fall within a range of $[a/1.5, a \times 1.5]$, where $a$ is the value of the third-last free parameter in MSMC, so the time segment just before the segments that are checked. If this condition is not fulfilled, we correct the estimates in the checked time intervals to $a$. This autocorrection is independently performed for each pair of haplotypes analysed (so for example we independently check $\lambda_{11}$, $\lambda_{12}$ and $\lambda_{22}$ independently).

We also constrain parameters $N_1(t)$ and $N_2(t)$ to be below $10^7$ and migration rates to be below 100, to avoid overflow issues during the fit. Furthermore, in MSMC-IM's automatic output report, we do not report estimated migration rates for times more ancient than after $M(t)$ has reached 0.999, because of the very little data that is left to infer migration rates when all but 0.1% of lineages have effectively already merged in one ancestral population.

##### 2.3.4 Interpreting Population size estimates

In MSMC-IM, we have two populations that never merge into one ancestral population. Instead, continuous migration is used to model movement of lineages across population boundaries, and hence also coalescence events between linegaes sampled across populations.

The degree to which lineages get mixed, looking back in time, can be quantified by the cumulative migration density, as defined in Methods as

$$M(t) = 1 - e^{-\int_0^t m(t')\mathrm{d}t'} \tag{64}$$

In recent times, where $M(t) \ll 1$, population sizes parameters $N_1(t)$ and $N_2(t)$ correspond closely to the inverse coalescence rates $1/\lambda_{11}(t)$ and $1/\lambda_{11}(t)$ estimtated by MSMC. However, as $M(t)$ approaches 1, the interpretation of these parameters differs from what one would normally call an "ancestral population size" in a clean-split model: In our model, we maintain two separate populations, so that with probability 1/2, two lineages will be in separate populations and cannot coalesce. Therefore, the effective coalescence rates in MSMC-IM for times at which $M(t) \to 1$, is half the rate expected for an ancestral population with size $N_1(t)$ or $N_2(t)$.

Therefore, for $M(t) \to 1$, a meaningful estimate for the effective "ancestral" population size would be $2N_1(t) \approx 2N_2(t)$. We therefore found it useful to report "corrected" population size estimates defined as

$$\begin{aligned} \mathbf{N_1'(t)} &= (1 - M(t))N_1(t) + M(t)2N_1(t) \\ \mathbf{N_2'(t)} &= (1 - M(t))N_2(t) + M(t)2N_2(t) \end{aligned} \tag{65}$$

### References


[1] Richard Durbin, Sean R Eddy, Anders Krogh, and Graeme Mitchison. *Biological sequence analysis: probabilistic models of proteins and nucleic acids.* Cambridge university press, 1998.

[2] Asger Hobolth, Lars Nørvang Andersen, and Thomas Mailund. On computing the coalescence time density in an isolation-with-migration model with few samples. *Genetics*, 187(4):1241–1243, April 2011.

[3] Heng Li and Richard Durbin. Inference of human population history from individual whole-genome sequences. *Nature*, 475(7357):493–496, July 2011.

[4] Anna-Sapfo Malaspinas, Michael C Westaway, Craig Muller, Vitor C Sousa, Oscar Lao, Isabel Alves, Anders Bergström, Georgios Athanasiadis, Jade Y Cheng, Jacob E Crawford, Tim H Heupink, Enrico Macholdt, Stephan Peischl, Simon Rasmussen, Stephan Schiffels, Sankar Subramanian, Joanne L Wright, Anders Albrechtsen, Chiara Barbieri, Isabelle Dupanloup, Anders

Eriksson, Ashot Margaryan, Ida Moltke, Irina Pugach, Thorfinn S Korneliussen, Ivan P Levkivskyi, J Víctor Moreno-Mayar, Shengyu Ni, Fernando Racimo, Martin Sikora, Yali Xue, Farhang A Aghakhanian, Nicolas Brucato, Søren Brunak, Paula F Campos, Warren Clark, Sturla Ellingvåg, Gudjugudju Fourmile, Pascale Gerbault, Darren Injie, George Koki, Matthew Leavesley, Betty Logan, Aubrey Lynch, Elizabeth A Matisoo-Smith, Peter J McAllister, Alexander J Mentzer, Mait Metspalu, Andrea B Migliano, Les Murgha, Maude E Phipps, William Pomat, Doc Reynolds, François-Xavier Ricaut, Peter Siba, Mark G Thomas, Thomas Wales, Colleen Ma'run Wall, Stephen J Oppenheimer, Chris Tyler-Smith, Richard Durbin, Joe Dortch, Andrea Manica, Mikkel H Schierup, Robert A Foley, Marta Mirazon Lahr, Claire Bowern, Jeffrey D Wall, Thomas Mailund, Mark Stoneking, Rasmus Nielsen, Manjinder S Sandhu, Laurent Excoffier, David M Lambert, and Eske Willerslev. A genomic history of aboriginal australia. *Nature*, September 2016.

[5] Paul Marjoram and Jeff D Wall. Fast "coalescent" simulation. *BMC Genet.*, 7:16, January 2006.

[6] Gilean A T McVean and Niall J Cardin. Approximating the coalescent with recombination. *Philos. Trans. R. Soc. Lond. B Biol. Sci.*, 360(1459):1387–1393, July 2005.

[7] Stephan Schiffels and Richard Durbin. Inferring human population size and separation history from multiple genome sequences. *Nat. Genet.*, 46(8):919–925, August 2014.
